# Acyl-CoA dehydrogenase substrate promiscuity limits the potential for development of substrate reduction therapy in disorders of valine and isoleucine metabolism

**DOI:** 10.1101/2022.11.22.517273

**Authors:** Sander M. Houten, Tetyana Dodatko, William Dwyer, Hongjie Chen, Brandon Stauffer, Robert J. DeVita, Frédéric M. Vaz, Chunli Yu, João Leandro

**Affiliations:** Department of Genetics and Genomic Sciences, Icahn School of Medicine at Mount Sinai, New York, NY 10029, USA; Mount Sinai Genomics, Inc, Stamford, CT 06902, USA; Department of Pharmacological Sciences, Icahn School of Medicine at Mount Sinai, New York, NY 10029, USA; Drug Discovery Institute, Icahn School of Medicine at Mount Sinai, New York, NY 10029, USA; Amsterdam UMC location University of Amsterdam, Department of Clinical Chemistry and Pediatrics, Laboratory Genetic Metabolic Diseases, Emma Children’s Hospital, Meibergdreef 9, Amsterdam, The Netherlands; Amsterdam Gastroenterology Endocrinology Metabolism, Inborn errors of metabolism, Amsterdam, The Netherlands; Core Facility Metabolomics, Amsterdam UMC location University of Amsterdam, Amsterdam, The Netherlands

## Abstract

Toxicity of accumulating substrates is a significant problem in several disorders of valine and isoleucine degradation notably short-chain enoyl-CoA hydratase (ECHS1 or crotonase) deficiency, 3-hydroxyisobutyryl-CoA hydrolase (HIBCH) deficiency, propionic acidemia (PA) and methylmalonic aciduria (MMA). Isobutyryl-CoA dehydrogenase (ACAD8) and short/branched-chain acyl-CoA dehydrogenase (SBCAD, *ACADSB*) function in the valine and isoleucine degradation pathways, respectively. Deficiencies of these acyl-CoA dehydrogenase (ACAD) enzymes are considered biochemical abnormalities with limited or no clinical consequences. We investigated whether substrate reduction therapy through inhibition of ACAD8 and SBCAD can limit the accumulation of toxic metabolic intermediates in disorders of valine and isoleucine metabolism. Using analysis of acylcarnitine isomers, we show that 2-methylenecyclopropaneacetic acid (MCPA) inhibited SBCAD, isovaleryl-CoA dehydrogenase, short-chain acyl-CoA dehydrogenase and medium-chain acyl-CoA dehydrogenase, but not ACAD8. MCPA treatment of wild-type and PA HEK-293 cells caused a pronounced decrease in C3-carnitine. Furthermore, deletion of *ACADSB* in HEK-293 cells led to an equally strong decrease in C3-carnitine when compared to wild-type cells. Deletion of *ECHS1* in HEK-293 cells caused a defect in lipoylation of the E2 component of the pyruvate dehydrogenase complex, which was not rescued by *ACAD8* deletion. MCPA was able to rescue lipoylation in *ECHS1* KO cells, but only in cells with prior *ACAD8* deletion. SBCAD was not the sole ACAD responsible for this compensation, which indicates substantial promiscuity of ACADs in HEK-293 cells for the isobutyryl-CoA substrate. Substrate promiscuity appeared less prominent for 2-methylbutyryl-CoA at least in HEK-293 cells. We suggest that pharmacological inhibition of SBCAD to treat PA should be investigated further.

## Introduction

Toxicity of accumulating substrates is a significant problem in many inborn errors of metabolism. Propionic acidemia (PA; MIM 606054) is caused by propionyl-CoA carboxylase deficiency and leads to toxic accumulation of propionyl-CoA and derivatives such as propionic, 3-hydroxypropionic and 2-methylcitric acid, as well as propionylcarnitine (C3-carnitine) and propionylglycine due to increased alternative metabolism (Ando et al 1972; Ando et al 1972; Weidman and Drysdale 1979; Kurczynski et al 1989). Methylmalonic aciduria due to methylmalonyl-CoA mutase deficiency or defects in cobalamin metabolism (MMA; MIM 251000, 251100, 251110) is characterized by similar metabolite perturbations, but with methylmalonic acid as the main accumulating metabolite. Valine and isoleucine degradation are thought to be the major source of propionyl-CoA and derived metabolites in these disorders (Thompson et al 1990; Leonard 1997). In addition, specific defects in valine metabolism are also characterized by pronounced toxicity of intermediates. Short-chain enoyl-CoA hydratase (ECHS1 or crotonase) deficiency (MIM 616277) and 3-hydroxyisobutyryl-CoA hydrolase (HIBCH) deficiency (MIM 250620) lead to the accumulation methacrylyl-CoA and acrylyl-CoA causing Leigh or Leigh-like syndrome (Loupatty et al 2007; Ferdinandusse et al 2013; Peters et al 2014; Peters et al 2015; Sakai et al 2015). Methacrylyl-CoA is a reactive intermediate that is thought to cause secondary defects in the respiratory chain and the pyruvate dehydrogenase complex (PDHc). Substrate reduction mostly through medical diets and emergency interventions is part of the treatment for many of these disorders. Specifically PA and MMA patients should limit intake of all propiogenic amino acids including valine and isoleucine (Manoli et al 1993; Baumgartner et al 2014; Fraser and Venditti 2016). Although mortality has declined with improved clinical management, the outcome of PA and MMA is still unfavorable as many patients have long-term complications that can affect multiple organ systems (de Baulny et al 2005; Touati et al 2006; Grunert et al 2012; Grunert et al 2013; Haijes et al 2019). In addition, medical foods resulted in iatrogenic amino acid deficiencies that were associated with adverse growth outcomes (Fraser and Venditti 2016; Manoli et al 2016). Therefore, there is a need for new therapeutic options such as pharmacological substrate reduction that may further improve outcome in PA. A low valine diet appeared beneficial in patients with ECHS1 and HIBCH deficiency, however, additional studies are needed to fully understand clinical outcomes (Soler-Alfonso et al 2015; Shayota et al 2019; Xu et al 2019; Abdenur et al 2020; Sato-Shirai et al 2021; Pata et al 2022). The degradation of all three branched-chain amino acids (BCAA; valine, leucine and isoleucine) starts with transamination and oxidative decarboxylation by shared enzymes (**Fig. S1**) (Neinast et al 2019). The third step of BCAA degradation is catalyzed by three specific acyl-CoA dehydrogenases (ACADs). Isobutyryl-CoA dehydrogenase (ACAD8) acts on isobutyryl-CoA, which is derived from valine, whereas short/branched-chain acyl-CoA dehydrogenase (SBCAD) is specific for the isoleucine degradation intermediate (*S*)-2-methylbutyryl-CoA. Inborn errors have been reported for both enzymes, namely isobutyryl-CoA dehydrogenase deficiency (MIM 611283) caused by mutations in *ACAD8* (Nguyen et al 2002), and 2-methylbutyrylglycinuria (MIM 610006) caused by mutations in *ACADSB* (Andresen et al 2000; Gibson et al 2000). Both deficiencies, however, are currently considered biochemical phenotypes of questionable clinical significance (Goodman and Duran 2014). Such conditions are diagnosed through biochemical and genetic methods, but are considered not harmful in humans. Most ACAD8 and SBCAD-deficient individuals diagnosed by newborn screening remain *asymptomatic* without treatment and low protein diet was without positive effects in *symptomatic* cases (Roe et al 1998; Andresen et al 2000; Gibson et al 2000; Sass et al 2004; Kanavin et al 2007; Oglesbee et al 2007; Sass et al 2008; Alfardan et al 2010; Pena et al 2012; Porta et al 2019; Eleftheriadou et al 2021; Feng et al 2021). Therefore, associations of ACAD8 and SBCAD deficiency with clinical disease are likely due to ascertainment bias. Based on this evidence from human genetics, ACAD8 and SBCAD are two putative candidates for safe pharmacological inhibition. Herein, we investigate whether substrate reduction therapy by inhibition of ACAD8 or SBCAD can limit the accumulation of toxic metabolic intermediates in PA, MMA and ECHS1 and HIBCH deficiencies.

## Material and Methods

### Cell lines

HEK-293 cells (ATCC, CRL-1573) were cultured in DMEM with 4.5 g/L glucose, 584 mg/L L-glutamine and 110 mg/L sodium pyruvate, supplemented with 10% fetal bovine serum (FBS), 100 U/mL penicillin, 100 μg/mL streptomycin, in a humidified atmosphere of 5% CO2 at 37ºC.

### Generation of CRISPR-Cas9 knockout cell lines

HEK-293 cell lines with a knockout (KO) for a gene of interest were generated using the CRISPR-Cas9 genome editing technique essentially as described (Ran et al 2013), with minor modifications (Violante et al 2019; Leandro et al 2020). For each gene, at least two different guides were chosen and cloned into the pSpCas9(BB)-2A-GFP vector (**Table S1**). Following plasmid purification, HEK-293 cells were transfected using Lipofectamine 2000. Twenty-four to 48 hours after transfection, cells with GFP signal were sorted as single cells into 96 well plates by FACS analysis. After expansion, the clonal cell lines were characterized by DNA sequencing (**Table S2**), immunoblotting and functional analysis essentially as described (Violante et al 2019; Leandro et al 2020). Since HEK-293 cell lines are hypotriploid, resolving the exact mutation for all each clonal cell lines is challenging, but all mutations were demonstrated to be deleterious through immunoblotting and functional assays.

### Immunoblotting

Cells were lysed in RIPA buffer supplemented with protease inhibitors cocktail (Roche), centrifuged 10 min at 12,000 xg, 4°C and total protein determined by the BCA method (Thermo Fisher Scientific). Proteins were separated on a Bolt 4–12% Bis-Tris Plus Gel (Thermo Fisher Scientific), blotted onto a nitrocellulose membrane (926-31092, LI-COR) and detected using the following primary antibodies: PCCB (PA5-57880; Invitrogen), SBCAD (HPA041458; Sigma-Aldrich), ACAD8 (HPA043903; Sigma-Aldrich), MCAD (55210-1-AP; Proteintech), lipoic acid (ab58724; Abcam, or 437695; Calbiochem), total OXPHOS human WB antibody cocktail (ab110411; Abcam) and citrate synthase (GT1761; GTX628143, Genetex). Proteins were visualized using IRDye 800CW or IRDye 680RD secondary antibodies (926-32210, 926-68070, 926-32211, 926-68071; LI-COR) in a Odyssey CLx Imager (LI-COR) with Image Studio Lite software (version 5.2, LI-COR). Equal loading was checked by Ponceau S (Acros Organics) staining and the citrate synthase signal.

### Metabolite analyses and MCPA studies

The acylcarnitine profiling in HEK-293 cells was performed essentially as previously described (Violante et al 2019). HEK-293 KO cells were seeded in 12-well plates and incubated overnight at 37°C. The following day, the media were removed, and 1 ml of minimal essential medium (MEM including 0.4 mM L-valine, 0.4 mM L-isoleucine and 0.4 mM L-leucine) supplemented with 0.4 mM L-carnitine (C0158, Sigma-Aldrich) was added. After incubation for 72 hours in a humidified CO2 incubator (5% CO2, 95% air) at 37°C, the medium was collected, and the cells were washed and resuspended in 200 μl of RIPA buffer to measure protein content using the BCA method and human serum albumin as standard.

For the characterization of 2-methylenecyclopropaneacetic acid (MCPA; M304360, Toronto Research Chemicals) in HEK-293 cells, the inhibitor was dissolved in anhydrous DMSO as a 20 mM stock concentration and stored at -20°C. The inhibitor was added to MEM at final concentrations of 0–200 μM, supplemented with 0.4 mM L-carnitine, 0.4% bovine serum albumin (fatty acid free) and 120 μM palmitic acid (C16:0). Acylcarnitines in media were measured by the Mount Sinai Biochemical Genetic Testing Lab (now Sema4). The carnitine ester of MCPA was observed as m/z 302.5, but not quantified. Acylcarnitine isomers were measured essentially as described (van Weeghel et al 2012) using the following transitions: m/z 246.2 > 85.0 for C5-carnitines and m/z 232.2 > 85.0 for C4-carnitines.

For the generation of samples for immunoblotting of lipoic acid, cell lines were treated with 25μM MCPA for 72 hours. The media included carnitine as described for the acylcarnitine profiling.

## Results

### Generation of HEK-293 cell line models for propionic acidemia and ECHS1 deficiency

We first generated HEK-293 cell line models for PA and ECHS1 deficiency using CRISPR-Cas9 genome editing to introduce null mutations in *PCCB* and *ECHS1*, respectively. Clonal *PCCB* and *ECHS1* KO cell lines were selected based on the presence of mutations in targeted gene, the absence of protein expression in an immunoblot and an expected biochemical abnormality in a functional test (**Fig. S2, S3**). Phenotypic variability between KO clones is a common challenge in genome editing experiments and may be related to off-target effects or heterogeneity of wild-type cells (Westermann et al 2022). In our experiments, we evaluated multiple clones in order to establish that the observed phenotype is reproducible.

For the PA cell line model, we measured extracellular C3-carnitine accumulation in four selected *PCCB* KO cell lines with absent PCCB upon immunoblotting (**Fig. S2A**). C3-carnitine levels were modestly increased (2.1-to 2.3-fold) in three out of the four *PCCB* KO cell lines when compared to the parental wild-type cells (**Fig. S2B**).

For the ECHS1 deficiency cell line model, we used an antibody against lipoic acid that detects lipoylated proteins such as the E2 component of the PDHc (E2p, DLAT) and the 2-oxoglutarate dehydrogenase complex (E2o, DLST). A specific decrease in the E2p has been observed in multiple tissues of patients with ECHS1 deficiency (Ferdinandusse et al 2015). We observed a 3.4-fold decrease in lipoylation signal for E2p and a 2.6-fold decrease in the lipoylation signal for E2o in all *ECHS1* KO cell lines when compared to controls (**Fig. S3A, B**). Activity of the 2-oxoglutarate dehydrogenase complex was 1.3-fold lower in *ECHS1* KO cell lines (**Fig. 3C**). By using an antibody cocktail against one subunit for each oxidative phosphorylation complex, we did not obtain evidence that the respiratory chain was affected in these *ECHS1* KO cell lines (not shown).

### MCPA inhibits SBCAD but not ACAD8 in HEK-293 cells

We next used hypoglycin A (2-methylenecyclopropanealanine, MCPA) as a tool compound for inhibition of multiple ACADs. MCPA inhibits SBCAD, isovaleryl-CoA dehydrogenase (IVD), short-chain acyl-CoA dehydrogenase (SCAD), medium-chain acyl-CoA dehydrogenase (MCAD) and glutaryl-CoA dehydrogenase (GCDH) (Tanaka et al 1971; Tanaka et al 1972; Hyman and Tanaka 1984), but not ACAD8 (Tanaka et al 1971; Ibrahim 2003; Ibrahim and Mohsen 2011). In order to characterize the metabolic effects of MCPA in HEK-293 cells, we analyzed acylcarnitine isomers in media to assess the degree of inhibition of the different ACAD enzymes.

C4-carnitine was separated in two isomers, isobutyryl-carnitine and butyryl-carnitine reflecting isobutyryl-CoA, the ACAD8 substrate, and butyryl-CoA, the SCAD substrate. MCPA decreased rather than increased isobutyryl-carnitine levels, indicating that MCPA does not inhibit ACAD8 (**Fig. 1**). Butyryl-carnitine was therefore solely responsible for the increase in C4-carnitine, and thus SCAD was inhibited by MCPA (IC50 = 1.6 ± 0.3 μM, **Fig. 1**). The level of butyryl-carnitine decreased at MCPA concentrations higher than 25 μM, which likely reflects a reduction in the supply of SCAD substrates due to the inhibition of MCAD, which functions upstream in the fatty acid β-oxidation pathway.

**Figure 1.**
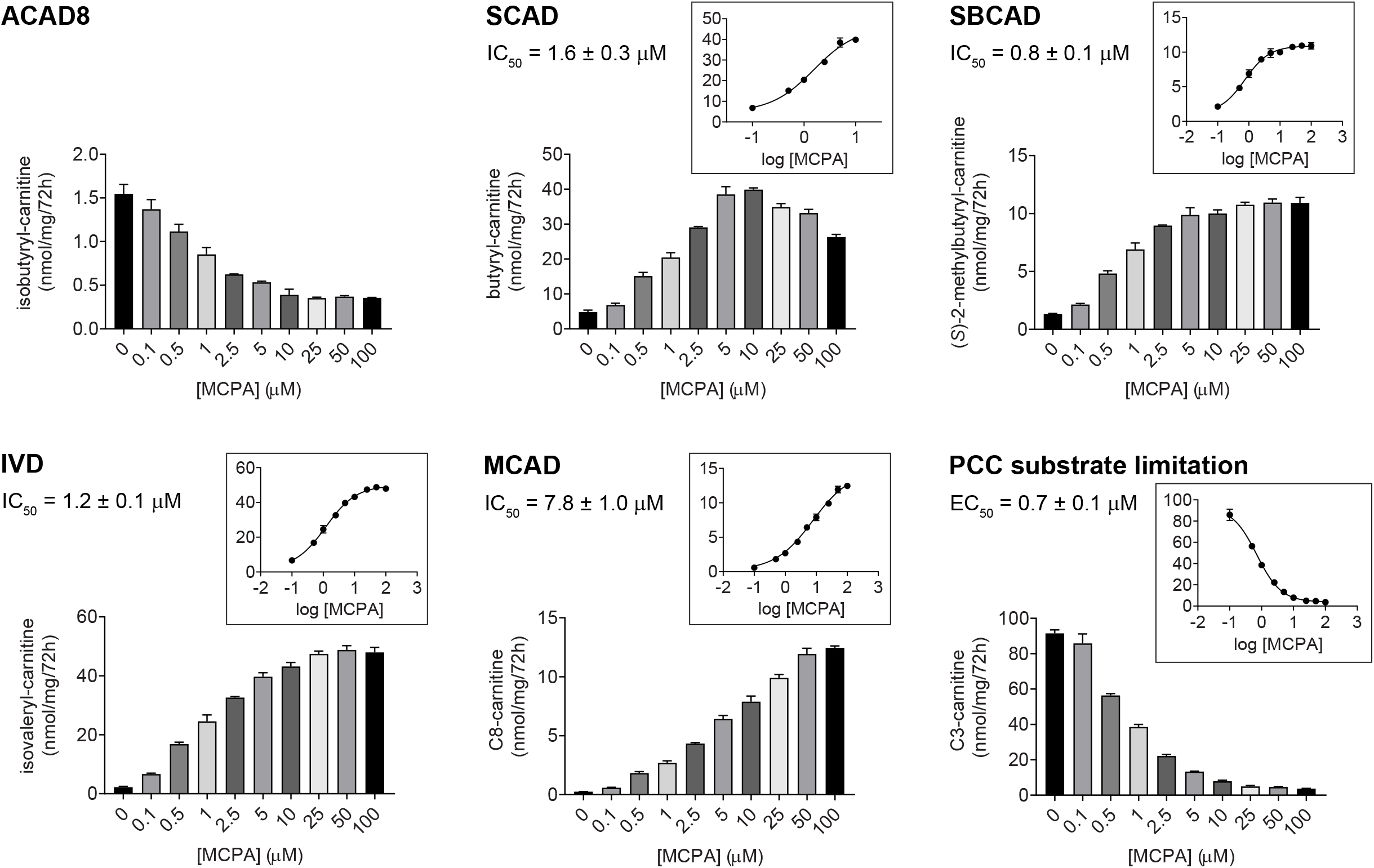
MCPA inhibits SBCAD, but not ACAD8, and lowers C3-carnitine. Quantification of C4- and C5-carnitine isomers, and C8-carnitine in the extracellular medium of HEK-293 cell lines in the presence of 0-100 μM MCPA. IC50 values for inhibition of SBCAD, IVD, MCAD were calculated using the full range of MCPA concentrations (0-100 μM). The IC50 value for inhibition of SCAD was calculated using MCPA concentrations up to 10 μM, at which the highest butyryl-carnitine concentration was reached. The estimated IC50 value did not change by much if we used concentration up to 5 (IC50 = 1.9 ± 0.9 μM) or 25 μM (IC50 = 1.1 ± 0.2 μM). C3-carnitine data were fitted to an EC50 curve demonstrating the dose dependent reduction of PCC substrate. The insets represent the corresponding IC50 or EC50. Error bars indicate SD.

C5-carnitine was separated in five isomers that included (*S*)-2-methylbutyryl-carnitine and isovaleryl-carnitine, surrogate markers of (*S*)-2-methylbutyryl-CoA and isovaleryl-CoA, the substrates of SBCAD and IVD, respectively. We observed an increase in (*S*)-2-methylbutyryl-carnitine levels indicating potent inhibition of SBCAD (IC50 = 0.8 ± 0.1 μM, **Fig. 1**). Isovaleryl-carnitine levels also increased revealing inhibition of IVD (IC50 = 1.2 ± 0.1 μM, **Fig. 1**).

C8-carnitine levels displayed a pronounced increase, but only at higher MCPA concentrations (IC50 = 7.8 ± 1.0 μM, **Fig. 1**). This indicated that the inhibition of MCAD by MCPA is not as potent when compared to the inhibition of various short-chain ACADs. No changes in C16-, C12- and C5DC-carnitine levels were detected (**Fig. S4**). Indeed, palmitoyl-CoA dehydrogenase (i.e. VLCAD and LCAD) is not inhibited by MCPA (Veitch et al 1987; Ikeda and Tanaka 1990). GCDH is known to be inhibited by MCPA (Hyman and Tanaka 1984), but in other studies we found that C5DC-carnitine accumulation due to GCDH deficiency is better detected in cell pellet extracts (Leandro et al 2020).

We noted a parallel decrease in C3-carnitine, which reflects the PCC substrate C3-CoA. (**Fig. 1**). This suggests that in HEK-293 cells, SBCAD contributes a significant portion of the production of C3-CoA. Indeed, the concentration of half-maximal response (EC50 = 0.7 ± 0.1 μM) closely reflects the IC50 of SBCAD inhibition by MCPA.

### ACADSB deletion limits the production of C3-carnitine in HEK-293 cells

We then explored further if inhibition of SBCAD can limit accumulation in C3-carnitine by generating *ACADSB* KO HEK-293 cell lines. Clonal *ACADSB* KO cell lines were selected based on the presence of *ACADSB* mutations and the absence of SBCAD protein in an immunoblot (**Fig. S5A**). As predicted, *ACADSB* KO cell lines caused a 10-fold (range 4 to 11-fold in individual clones) increase in C5-carnitine levels (**Fig. 2A and Fig. S5B**). The average decrease of C3-carnitine was 4-fold (range 2.6 to 7.9-fold in individual clones) in *ACADSB* KO cell lines when compared to the wild type HEK-293 cell line (**Fig. 2A and Fig. S5A**). Smaller changes were observed in some other acylcarnitines notably C4-, C4OH-and C5:1-carnitine (**Fig. S5C**). In contrast, the decrease in C3-carnitine was only 1.5-fold (range 1.2 to 1.8-fold in individual clones) in *ACAD8* KO cell lines when compared to the wild-type HEK-293 cell line (**Fig. S7**). This further confirms a more prominent role of SBCAD in the production of C3-carnitine in HEK-293 cells.

**Figure 2.**
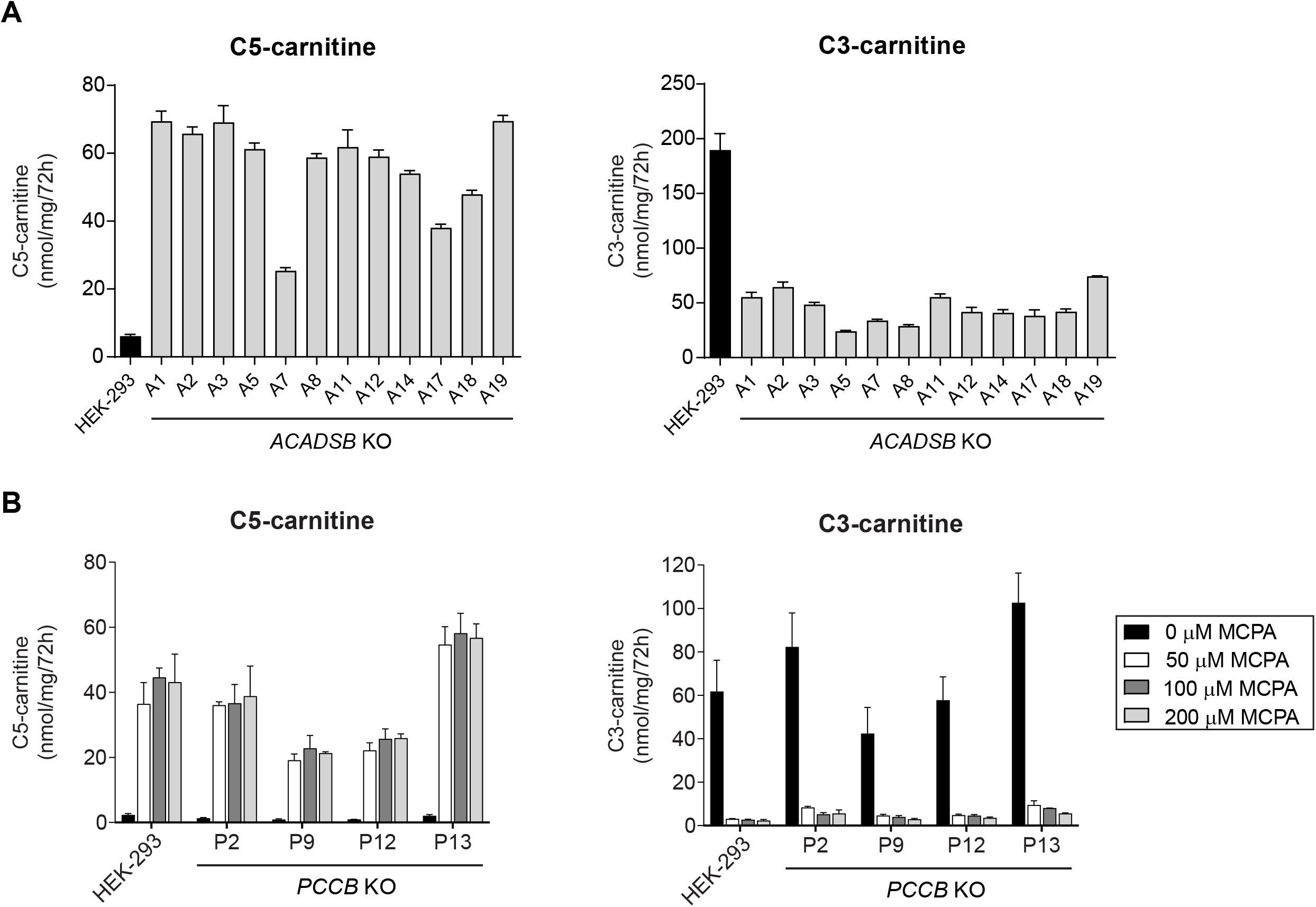
Deletion of *ACADSB* in HEK-293 cells increases C5-carnitine and limits the accumulation C3-carnitine reflecting PCC substrate limitation. (A) Production of C5-carnitine and C3-carnitine in the extracellular medium of selected wild-type and *ACADSB* KO HEK-293 cell lines. Error bars indicate SD. (B) Production of C5- and C3-carnitine in the extracellular medium of selected wild-type and *PCCB* KO HEK-293 cell lines in the presence of increasing concentrations of the ACAD inhibitor MCPA.

**Figure 3.**
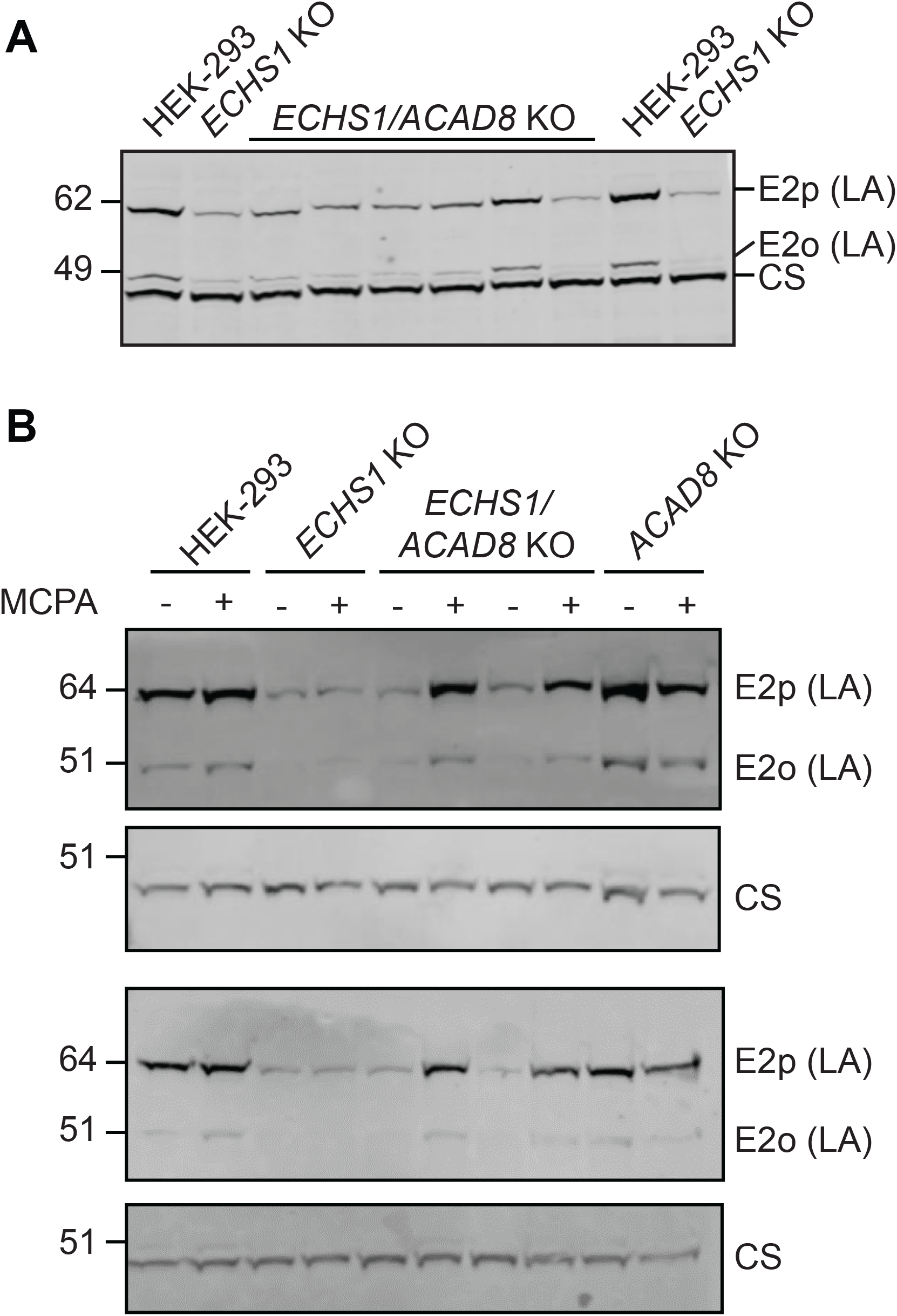
Rescue of lipoylation by MCPA in *ECHS1*/*ACAD8* KO cell lines. (A) Lipoylation of E2p and E2o in *ECHS1*/*ACAD8* KO cell lines. Western blots of cell lysates were probed with an antibody against lipoic acid, and E2p (DLAT) and E2o (DLST) were identified based on their molecular weight. Citrate synthase was used as loading control and the position of the molecular weight marker proteins is indicated. (B) *ECHS1*/*ACAD8* KO cell lines and selected other cell lines were treated with 25μM MCPA or vehicle for 72 hours as indicated.

In order to further establish that inhibition of SBCAD can limit the metabolite accumulation in PA, we again used MCPA. We measured C3-carnitine levels in wild-type and four *PCCB* KO cell lines (**Fig. 2B**). A pronounced reduction in C3-carnitine was observed for both wild-type (27-fold) and *PCCB* KO cell lines (between 15 to 19-fold). Corresponding increases in the levels of C5-carnitine (**Fig. 2B**), as well as other short- and medium-chain acylcarnitines (C4-, C6- and C8-carnitine) (**Fig. S6**), indicated successful inhibition of the ACAD enzymes.

### ACAD8 KO is necessary, but not sufficient to rescue decreased lipoylation in ECHS1 deficiency

ACAD8 is thought to be the primary source of the toxic valine-derived methacrylyl-CoA that decreases lipoylation of E2p (DLAT) and E2o (DLST). To validate if ACAD8 could be a pharmacological target for the treatment of ECHS1 deficiency, we generated *ECHS1*/*ACAD8* double KO as well as *ACAD8* single KO HEK-293 cell lines. Since we were unable to detect ACAD8 protein in HEK-293 cells using a validated ACAD8 antibody, we selected *ACAD8* KO clones based on DNA sequencing results only. Most often the selected clones had small 1 or 2 nucleotide insertions or deletions leading to a frameshift and the introduction of a premature stop codon in combination with a large insertion (ranging from ∼90 to ∼270 basepairs) on the other allele. All selected *ACAD8* and *ECHS1*/*ACAD8* KO clones had C4-carnitine levels higher than wild-type HEK-293 cells (**Fig. S7**).

We next studied lipoylation in the *ECHS1*/*ACAD8* KO clones. Unexpectedly, *ACAD8* KO did not rescue the decreased lipoylation of E2p and E2o in the *ECHS1* KO cell line (**Fig. 3A**). This suggests that ACAD8 is either not involved in the production of the toxic intermediate, or its deficiency is compensated for by other ACADs with overlapping substrate specificity (i.e. substrate promiscuity). Given the firmly established role of ACAD8 in valine metabolism, the latter option is more likely. Therefore, we treated HEK-293, *ECHS1, ACAD8* and *ECHS1*/*ACAD8* KO cell lines with MCPA, which will inhibit SBCAD, IVD, SCAD, MCAD and GCDH. MCPA treatment rescued lipoylation of E2p and E2o in *ECHS1*/*ACAD8* KO cell lines, but not in *ECHS1* KO cell lines (**Fig. 3B**). These data demonstrate that although *ACAD8* KO is not sufficient in itself to rescue decreased lipoylation in ECHS1 KO cell lines, it is essential for rescue by MCPA. This strongly suggests that in addition to ACAD8, inhibition of one or more of the ACADs targeted by MCPA is necessary to rescue lipoylation in *ECHS1* KO cell lines.

### SBCAD is not the only ACAD able to compensate for loss of ACAD8 function

There is significant overlap in the *in vitro* substrate specificity of ACAD8 and SBCAD, which both accept 2-methyl short/branched-chain acyl-CoAs including isobutyryl-CoA and 2-methylbutyryl-CoA (Ikeda et al 1983; Ikeda and Tanaka 1983; Rozen et al 1994; Willard et al 1996; He et al 2003). Therefore, SBCAD is the most likely candidate ACAD able to compensate for ACAD8 deficiency. To test if combined inhibition of ACAD8 and SBCAD is sufficient to rescue deficient lipoylation in *ECHS1* KO cell lines, we generated *ECHS1*/*ACAD8*/*ACADSB* triple KO cell lines (**Fig. 4A**). The successful generation of *ECHS1*/*ACAD8*/*ACADSB* KO cell lines was demonstrated by the absence of SBCAD protein in an immunoblot (**Fig. 4A**) and increased C5-carnitine (**Fig. 4A, S8**). *ACADSB* KO did not rescue the decreased lipoylation of E2p in an *ECHS1*/*ACAD8* KO cell line (**Fig. 4A**). Treatment of these *ECHS1*/*ACAD8*/*ACADSB* KO cell lines with MCPA rescued lipoylation of E2p (**Fig. 4B**). This result suggests SBCAD is not the compensating ACAD, or it is not the only ACAD that is able to compensate for ACAD8 inhibition.

**Figure 4.**
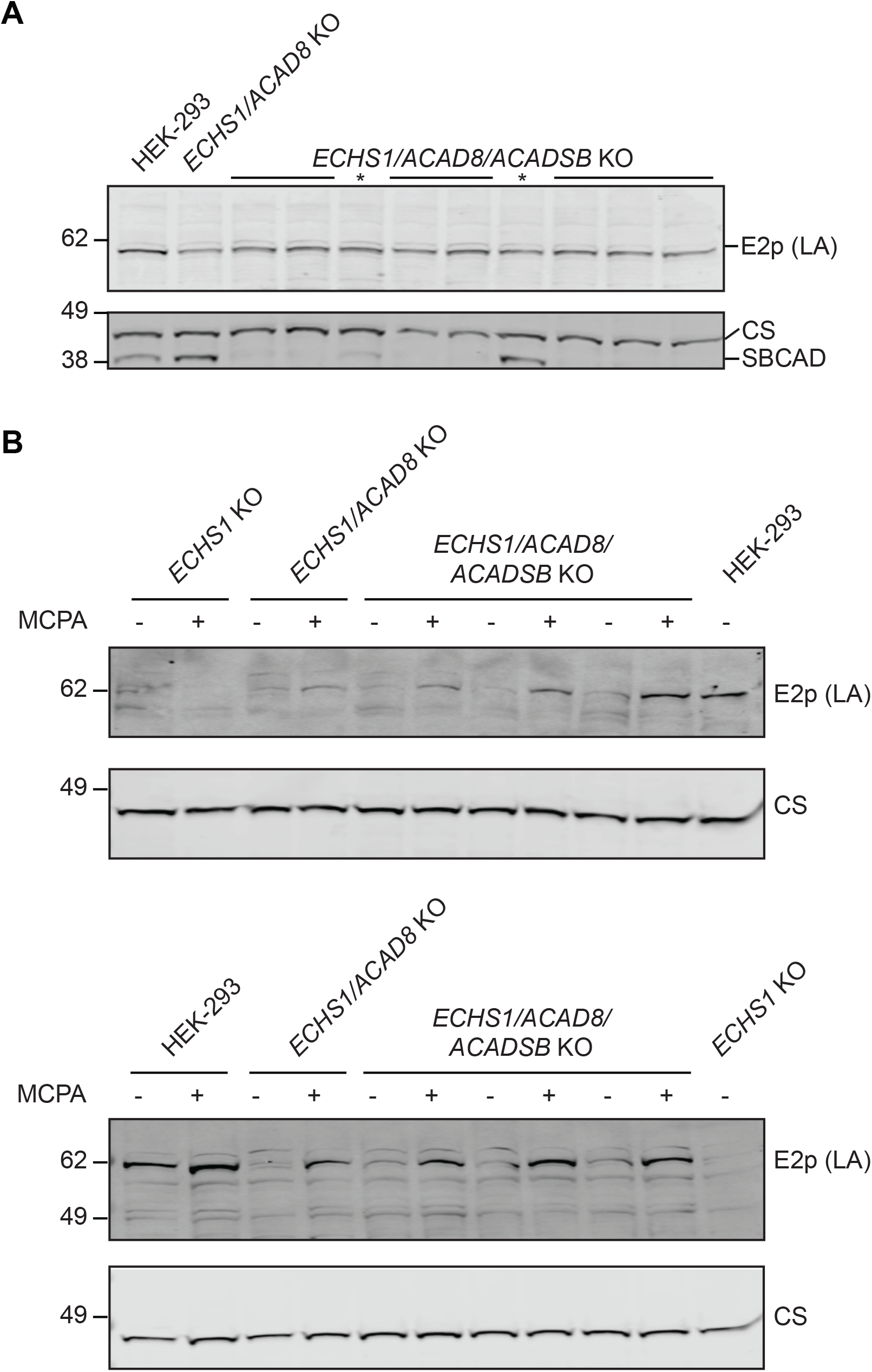
*ACADSB* KO is not sufficient to rescue of lipoylation in *ECHS1*/*ACAD8* KO cell lines. (A) Lipoylation of E2p in *ECHS1*/*ACAD8/ACADSB* KO cell lines. Western blots of cell lysates were probed with an antibody against lipoic acid and E2p (DLAT) was identified based on its molecular weight. E2o (DLST) was not reliably detected in these blots. Citrate synthase was used as loading control and the position of the molecular weight marker proteins is indicated. Two cell lines without incomplete *ACADSB* KO are marked with an asterisk. (B) *ECHS1*/*ACAD8*/*ACADSB* KO and selected other cell lines were treated with 25μM MCPA or vehicle for 72 hours as indicated.

### MCAD is not able to compensate for loss of ACAD8 and SBCAD

Recently, substrate promiscuity between ACAD8, SBCAD and MCAD was demonstrated on the flux of metabolites through the BCAA metabolic pathways in primary hepatocytes isolated from PA patients (Collado et al 2022). These results are reminiscent of our findings on the rescue of the lipoylation defect in ECHS1 deficient HEK-293 cells. Based on this report, we prioritized MCAD as another candidate over the other potential candidates SCAD and GCDH. We then generated *ECHS1*/*ACAD8*/*ACADSB/ACADM* quadruple and *ECHS1*/*ACAD8*/*ACADM* triple KO cell lines. The successful KO of *ACADM* was demonstrated by deficient MCAD protein in immunoblot and accumulation of C10-, C8- and C6-carnitine in the media (**Fig. S9**). The additional KO of *ACADM* did not lead to a clear rescue lipoylation of E2p (**Fig. 5A**). Treatment of these *ECHS1*/*ACAD8*/*ACADSB/ACADM* and *ECHS1*/*ACAD8*/*ACADM* KO cell lines with MCPA did rescue lipoylation of E2p to levels observed in wild-type HEK-293 cells (**Fig. 5B**). Similarly, treatment of *ECHS1*/*ACAD8*/*ACADSB/ACADM* and *ECHS1*/*ACAD8*/*ACADM* KO cell lines with MCPA rescued lipoylation of E2o (**Fig. 5B**).

**Figure 5.**
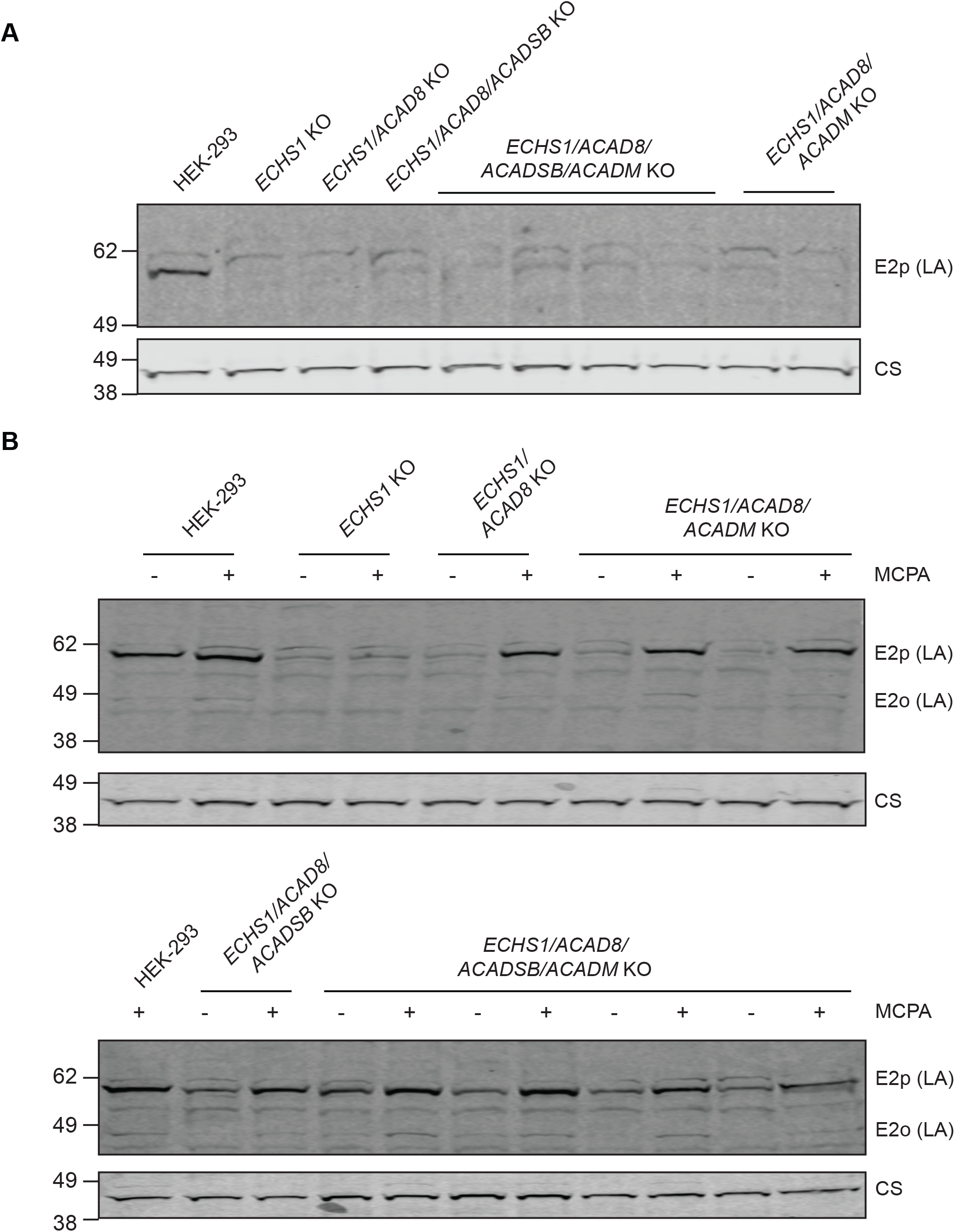
*ACADM* KO is not sufficient to rescue of lipoylation in *ECHS1*/*ACAD8/ACADSB* KO cell lines. (A) Lipoylation of E2p in *ECHS1*/*ACAD8/ACADSB/ACADM* KO cell lines. Western blots of cell lysates were probed with an antibody against lipoic acid and E2p (DLAT) was identified based on its molecular weight. E2o (DLST) was not reliably detected in these blots. Citrate synthase was used as loading control and the position of the molecular weight marker proteins is indicated. Two cell lines without incomplete *ACADSB* KO are marked with an asterisk. (B) *ECHS1*/*ACAD8*/*ACADSB/ACADM* KO and selected other cell lines were treated with 25μM MCPA or vehicle for 72 hours as indicated. A faint signal for E2o was noted in these immunoblots.

## Discussion

Inborn errors of valine and isoleucine metabolism are characterized by significant toxicity of accumulating substrates. Substrate reduction through metabolic maintenance and emergency treatment is an important pillar of the treatment of these disorders. We used pharmacological (MCPA) and genetic (CRISPR-Cas9 genome editing) tools in the HEK-293 cell line to investigate if pharmacological inhibition of SBCAD and ACAD8 may be a useful alternative approach for substrate reduction. We found that SBCAD is responsible for the majority of C3-carnitine production and found only a minor contribution of ACAD8. We speculate that inhibition of SBCAD should be further investigated as a novel therapeutic in PA and MMA. We also observed that loss-of-ACAD8 function was unable to rescue deficient lipoylation of E2p and E2o in ECHS1 deficiency. By using MCPA, we found that in addition to ACAD8, inhibition of other ACADs was necessary to fully rescue the deficient lipoylation, but were unable to establish the identity of these ACADs with the current results. Our work is consistent with substrate promiscuity of ACAD8, SBCAD and other ACADs most notably for the isobutyryl-CoA substrate (Ikeda et al 1983; Ikeda and Tanaka 1983; Rozen et al 1994; Willard et al 1996; He et al 2003; Collado et al 2022). Substrate promiscuity of enzymes poses challenges for the development of substrate reduction therapy, not only in disorders of valine and isoleucine metabolism, but also in glutaric aciduria type 1 (Leandro et al 2020).

The primary roles of ACAD8 and SBCAD in valine and isoleucine metabolism, respectively, are firmly established by the characterization of accumulating metabolites in individuals with ACAD8 and SBCAD deficiencies. ACAD8 deficiency leads to isobutyrylglycinuria and increased isobutyrylcarnitine (C4-carnitine). SBCAD deficiency leads to 2-methylbutyrylglycinuria and elevated (*S*)-2-methylbutyrylcarnitine (C5-carnitine). Importantly, there is no apparent C5-carnitine accumulation in ACAD8 deficiency, and no C4-carnitine accumulation in SBCAD deficiency. There are indeed significant *in vitro* substrate promiscuities of ACAD8 and SBCAD, both of which use 2-methyl short/branched-chain acyl-CoAs including isobutyryl-CoA and 2-methylbutyryl-CoA. In fact, based on biochemical studies, SBCAD was initially thought to be involved in both isoleucine and valine metabolism (Ikeda et al 1983; Ikeda and Tanaka 1983). Accordingly, human and rat SBCAD are active with 2-methylbutyryl-CoA and isobutyryl-CoA as well as hexanoyl-CoA, but the highest catalytic efficiency is observed with 2-methylbutyryl-CoA (Ikeda and Tanaka 1983; Rozen et al 1994; Willard et al 1996; He et al 2003). Similarly, ACAD8 displays activity with isobutyryl-CoA and 2-methylbutyryl-CoA, but the highest catalytic efficiency is observed with the former (Ibrahim 2003; Ibrahim and Mohsen 2011). These observations, however, do not provide insight into a potential compensatory mechanism when either of these enzymes is defective. Such compensatory mechanism will likely depend on at least two factors. The first is mRNA/protein expression level with higher levels likely leading to a higher ability to compensate. The second factor is the concentration of the accumulating substrate that is needed to overcome the lower catalytic efficiency of the compensating ACAD. Previous work in ACAD8 and SBCAD-deficient fibroblasts and hepatocytes from PA patients treated with potent inhibitors of various ACAD enzymes indeed suggests that such compensatory mechanisms exist (McCalley et al 2019; Collado et al 2022). This has led these investigators to speculate that ACAD8 and SBCAD deficiencies are without clinical consequences because of this proposed enzymatic compensation (McCalley et al 2019; Collado et al 2022). It remains unknown whether simultaneous inhibition of ACAD8 and SBCAD is safe. Further studies in animal models such as *Acad8, Acadsb* and *Acad8*/*Acadsb* KO mice are warranted.

Our results show that inhibition of SBCAD by MCPA in wild-type HEK-293 cells and a cell line model of PA (*PCCB* KO) decreases C3-carnitine production, which indicates decreased cellular propionyl-CoA levels. A decrease in propionyl-CoA due to CoA sequestration is unlikely given that the levels of C2-carnitine displayed a less pronounced decrease. Moreover, the IC50 for SBCAD is virtually identical to the EC50 for C3-carnitine decrease further suggesting causality. In accordance with this major role of SBCAD, McCalley et al have demonstrated that isoleucine was the major amino acid contributing to C3-carnitine production in fibroblasts, whereas valine, methionine and threonine were less contributory (McCalley et al 2019). We next generated *ACADSB* KO HEK-293 cell lines to provide additional data to support that blocking the isoleucine degradation pathway at the level of SBCAD is sufficient to limit the accumulation of propionyl-CoA derivatives. We found that C5-carnitine was increased in the *ACADSB* KO cell lines, whereas C3-carnitine was decreased 4.2-fold. In contrast, C3-carnitine decreased only 1.5-fold in *ACAD8* KO cell lines. These data demonstrate that isoleucine is the major propiogenic precursor in HEK-293 cells. *In vivo* isotope tracing in mice showed that both valine and isoleucine are oxidized in all tissues analyzed including liver and brain (Neinast et al 2019) and thus contribute to propionyl-CoA production. Overall, amino acid degradation is thought to contribute ∼50% of whole body propionic acid production, whereas oxidation of odd-chain fatty acids and production by the gut microbiome are each estimated to contribute 25% (Thompson et al 1990; Leonard 1997). It is conceivable that despite the fact that isoleucine is not the only source of propionyl-CoA, SBCAD inhibition may be sufficient to achieve a therapeutic benefit in PA patients. The contribution of individual amino acids such as isoleucine to propionic acid production is unknown and will be addressed. We suggest the study of *Acad8* and *Acadsb* KO mouse models, which are available through the International Mouse Phenotyping Consortium (Groza et al 2022).

Along similar lines, we investigated if ACAD8 inhibition could rescue metabolite toxicity associated with ECHS1 deficiency. Our *ECHS1* KO HEK-293 cells showed a pronounced defect in lipoylation of E2p and E2o. A specific decrease in the E2p has been observed in multiple tissues of patients with ECHS1 deficiency (Ferdinandusse et al 2015), but lipoylation has not been studied yet. Methacrylyl-CoA and acryloyl-CoA are known to react with the thiol group of cysteine and cysteamine in a Michael-type addition. It is unknown if they react with thiol groups of dihydrolipoic acid. We observed that *ACAD8* KO could not rescue the deficient lipoylation in *ECHS1* KO cells. MCPA treatment rescued lipoylation in *ECHS1*/*ACAD8* DKO cell lines, but not in *ECHS1* single KO cell lines. This is consistent with the observation that MCPA does not inhibit ACAD8 and likely illustrates promiscuity for the isobutyryl-CoA substrate with other ACADs able to compensate for ACAD8 deficiency. Remarkably, we were unable to identify these other ACADs since sequential KO of the primary candidates *ACADSB* and *ACADM* was unsuccessful in rescuing deficient lipoylation. This is unambiguously illustrated by the observation that *ECHS1*/*ACAD8*/*ACADSB*/*ACADM* quadruple KO cell lines were still rescued by MCPA addition. It could be that substrate promiscuity of SCAD or GCDH with isobutyryl-CoA produces levels of methacrylyl-CoA that are high enough for toxicity especially given the relatively high concentration of valine in the cell culture media (0.4 mM). Although unlikely, it remains also possible that MCPA acts via another mechanism, i.e. not via ACAD inhibition. In addition to methacrylyl-CoA, acryloyl-CoA is a reported toxic metabolite in ECHS1 deficiency (Peters et al 2014). Acryloyl-CoA is derived from propionyl-CoA and thus in our cell line system its generation should largely be blocked by the KO of *ACADSB* and *ACAD8*. In addition, ECHS1 itself is involved in the generation of propionyl-CoA from valine and isoleucine, which is illustrated by the very low C3-carnitine levels upon *ECHS1* deletion (**Fig. S7 and S8**).

In summary, our data provide preliminary evidence that deletion of SBCAD can reduce accumulation of propionyl-CoA and its derivatives. As a result, it provides validation that pharmacological inhibition of SBCAD may provide benefit for patients with PA or other inborn errors of propionic acid and cobalamin metabolism.

## Supporting information

All supplemental documents

## Supplemental Information

Supplemental information is available online.

## Acknowledgments

We thank Henk van Lenthe of the Laboratory Genetic Metabolic Diseases, Amsterdam UMC for the analysis of acylcarnitine isomers. Research reported in this publication was supported by the Propionic Acidemia Foundation (PAF, initial research grant) and the Eunice Kennedy Shriver National Institute Of Child Health & Human Development of the National Institutes of Health under Award Number R03HD105916. The content is solely the responsibility of the authors and does not necessarily represent the official views of the National Institutes of Health.

## Conflict of Interest Statement

The authors declare no competing interests.

## Author contributions

Conception and design of the work described, SMH; Acquisition of data: TD, WD, JL; Analysis and interpretation of data: SMH, WD, HC, RJDV, BS, RS, FMV, CY, JL; Reporting of the work described: SMH, JL.

## Notes

### Competing Interest Statement

The authors have declared no competing interest.

