## Supplementary material for "Acyl-CoA dehydrogenase substrate promiscuity limits the potential for development of substrate reduction therapy in disorders of valine and isoleucine metabolism": All supplemental documents

#### Supplemental Figures

**Figure S1. The degradation pathway of branched-chain amino acids (BCAAs) and propionyl-CoA metabolism.** BCAAs undergo an initial reversible transamination step by the branched-chain aminotransferase isozymes BCAT1 (cytosolic) or BCAT2 (mitochondrial) to produce the corresponding 2-oxo acid analogs. The second enzymatic step proceeds via an irreversible oxidative decarboxylation and coupled thioesterification of the respective 2-oxo acids by the branched-chain 2-oxo acid dehydrogenase complex (BCKDHc). The BCKDHc is a multisubunit enzymatic complex comprising several copies of an E1 component, a heterotetramer of BCKDHA and BCKDHB, a thiamine diphosphate-dependent decarboxylase; an E2 component DBT, a lipoate-dependent transacylase, and an E3 component DLD, a dehydrogenase that is shared with the other 2-oxo acid dehydrogenase complexes; the pyruvate, 2-oxoglutarate and 2-oxoadipate dehydrogenase complexes. The BCKDHc kinase (BCKDK) phosphorylates serine 293 of the E1$\alpha$ subunit, which inactivates the BCKDHc. The BCKDHc is dephosphorylated by the mitochondrial protein phosphatase PPM1K. The three BCAAs share the BCAT and BCKDH steps, after which unique enzymes participate in the catabolism of each BCAA, with the exception of the hydratase step in valine and isoleucine metabolism catalyzed by crotonase (ECHS1). Valine is a gluconeogenic amino acid, producing succinyl-CoA, whereas leucine is ketogenic, producing acetoacetic acid and acetyl-CoA. Isoleucine is both ketogenic and gluconeogenic with the formation of acetyl-CoA and succinyl-CoA. In patients with mutations in the *PCCA* or *PCCB* gene, propionyl-CoA carboxylase (PCC) is deficient leading to accumulation of propionyl-CoA and its toxic derivatives (2-methylcitric acid, propionic acid, 3-OH-propionic acid, propionylglycine and propionyl-carnitine) causing propionic acidemia. Threonine and methionine catabolism, the oxidation of branched-chain and odd-chain fatty acids, and cholesterol side-chain metabolism also contribute to the pool of propionyl-CoA. The abbreviations used are: ACAD8, isobutyryl-CoA dehydrogenase (also known as acyl-CoA dehydrogenase family member 8); SBCAD, short/branched-chain acyl-CoA dehydrogenase (also 2-methylbutyryl-CoA dehydrogenase); ACAT1, acetyl-CoA acetyltransferase 1; AdoCbl, adenosylcobalamin; ALDH6A1, methylmalonate semialdehyde dehydrogenase; AUH, 3-methylglutaconyl-CoA hydratase; BCAT, branched-chain aminotransferase; BCKDH, branched-chain 2-oxo acid dehydrogenase; DBT, dihydrolipoamide branched-chain transacylase; DLD, dihydrolipoyl dehydrogenase; ECHS1, mitochondrial short-chain enoyl-CoA hydratase 1; HIBADH, 3-hydroxyisobutyric acid dehydrogenase; HIBCH, 3-hydroxyisobutyryl-CoA hydrolase; HMGCL, 3-hydroxy-3-methylglutaryl CoA lyase; HSD17B10,

2-methyl-3-hydroxybutyryl-CoA dehydrogenase (also known as 3-hydroxyacyl-CoA dehydrogenase type-2); IVD, isovaleryl-CoA dehydrogenase; MCCC1/2, methylcrotonyl-CoA carboxylase subunit $\alpha/\beta$; MCEE, methylmalonyl-CoA epimerase; MUT, methylmalonyl-CoA mutase; PCCA/B, propionyl-CoA carboxylase subunit $\alpha/\beta$.

**Figure S2. A HEK-293 cell line model for PA.** (A) Immunoblot validation of *PCCB* KO cell lines. *PCCB* KO cell lines were generated in wild type HEK-293 cells using guides targeting exon 5 (*upper panel*) or exon 7 (*lower panel*). Individual *PCCB* KO clones are indicated as P#. Citrate synthase (CS) was used as loading control and the position of molecular mass marker proteins (in kDa) is given. Selected clones are indicated in bold. (B) Production of C3-carnitine in the extracellular medium of selected *PCCB* KO cell lines (△) and wild-type HEK-293 cells (●). Error bars indicate SD.

**Figure S3. Characterization of *ECHS1* KO cell lines.** (A) *ECHS1* was targeted using CRISPR-Cas9 genome editing with guides in exon 2 and exon 3. For each guide, 10 clonal cell lines were characterized using immunoblotting. Wild-type HEK-293 cells served as control. For both guides, 8 out of 10 clones showed a complete loss of ECHS1 protein. The same blots were probed with an antibody against lipoic acid, and E2p (DLAT) and E2o (DLST) were identified based on their molecular weight. Citrate synthase (CS) was used as loading control and the position of the molecular weight marker proteins is indicated. (B) Quantification of E2p and E2o normalized to CS. (C) Activity of the 2-oxoglutarate dehydrogenase complex in wild-type HEK-293 and five selected *ECHS1* KO cell lines. Error bars indicate SD. ****, $P < 0.0001$.

**Figure S4. Acylcarnitine levels in HEK-293 cells after incubation with 0-100 μM MCPA (Related to Figure 1).** (A) Production of acylcarnitines in the extracellular medium of HEK-293 cells after incubation with 0-100 μM MCPA. (B) Quantification of C4- and C5-carnitine isomer levels upon inhibition of HEK-293 cells with MCPA. The fifth C5-carnitine isomer, pivaloyl-carnitine was below the level of detection. Error bars indicate SD.

**Figure S5. Validation of *ACADSB* KO cell lines (Related to Figure 2A).** (A) *ACADSB* KO cell lines were generated in wild type HEK-293 cells, using a guide targeting exon 6 5' strand (*upper panel*) or 3' strand (*lower panel*). *ACADSB* KO clones are indicated as A#. Citrate synthase (CS) was used as loading control and the position of molecular mass marker proteins (in kDa) is given. Selected clones are indicated in bold. (B) Statistical analyses of C3-carnitine and C5-

carnitine extracellular levels in wild type (●) and *ACADSB* KO (△) HEK-293 cell lines. There is a 4.2-fold reduction in C3-carnitine levels and a 9.3-fold increase in C5-carnitine levels in *ACADSB* KO cell lines compared with wild type cells. Error bars indicate SD. ****, P < 0.0001. (C) Production of acylcarnitines in the extracellular medium of wild type and selected *ACADSB* KO HEK-293 cell lines.

**Figure S6. Acylcarnitine levels in wild-type and *PCCB* KO HEK-293 cell lines after incubation with MCPA (Related to Figure 2B).** Production of acylcarnitines in the extracellular medium of wild-type and selected *PCCB* KO HEK-293 cell lines after incubation with increasing concentrations of MCPA. Error bars indicate SD.

**Figure S7. Validation of *ACAD8* KO cell lines (Related to Figure 3).** (A) Acylcarnitine levels in wild-type, ECHS1, ACAD8 and ECHS1/ACAD8 KO HEK-293 cell lines. Four independently generated clones were tested for the ACAD8 and ECHS1/ACAD8 KO HEK-293 cell lines. (B) Acylcarnitine levels in wild-type, ECHS1, ACAD8 and ECHS1/ACAD8 KO HEK-293 cell lines upon treatment with MCPA or vehicle (DMSO). Cell lines were treated with 25μM MCPA for 72 hours as indicated. The values for the *ACAD8* and *ECHS1*/*ACAD8* KO HEK-293 cell lines were the average of 4 different clones.

**Figure S8. Validation of *ECHS1*/*ACAD8*/*ACADSB* KO cell lines (Related to Figure 4).** (A) Acylcarnitine levels in wild-type, *ECHS1*, *ECHS1*/*ACAD8*, *ACAD8*/*ACADSB*, *ECHS1*/*ACADSB*, *ECHS1*/*ACAD8*/*ACADSB* KO HEK-293 cell lines. (B) Acylcarnitine levels in wild-type, *ECHS1*, *ECHS1*/*ACAD8*, *ECHS1*/*ACAD8*/*ACADSB* KO HEK-293 cell lines upon treatment with MCPA or vehicle (DMSO). Cell lines were treated with 25μM MCPA for 72 hours as indicated. The values for the *ECHS1*/*ACAD8*/*ACADSB* KO HEK-293 cell lines were the average of 3 different clones. Duplicate measures are plotted for wild-type HEK-293 and a single clone of the *ECHS1* and *ECHS1*/*ACAD8* cell lines.

**Figure S9. *ECHS1*/*ACAD8*/*ACADSB*/*ACADM* KO cell lines (Related to Figure 5).** (A) Immunoblot validation of *ACADM* KO cell lines obtained using a guide targeting exon 5. The position of the molecular weight marker proteins is indicated. Similar results were obtained for a guide targeting exon 7. Lipoylation was studied in clones obtained using guides targeting exon 5 and 7. (B) Acylcarnitine levels in wild-type, *ECHS1*/*ACAD8*, *ECHS1*/*ACAD8*/*ACADSB*, *ECHS1*/*ACAD8*/*ACADM* and *ECHS1*/*ACAD8*/*ACADSB*/*ACADM* KO HEK-293 cell lines upon treatment with 100 μM lauric acid (C12:0) or vehicle (ethanol). Cell lines were treated with 25μM MCPA for 72 hours as indicated. The values for the *ECHS1*/*ACAD8*/*ACADM* and *ECHS1*/*ACAD8*/*ACADSB*/*ACADM* KO HEK-293 cell lines were the average of 4 different clones. Duplicate measures are plotted for wild-type HEK-293 and a single clone of the *ECHS1*/*ACAD8* and *ECHS1*/*ACAD8*/*ACADSB* KO cell lines.

**Supplemental tables**

**Table S1. Guide sequences used for CRISPR-Cas9 genome editing.**

| Target gene | Exon | Guide sequence |
|---|---|---|
| *PCCB* | 5 (+ strand) | CATGGACCAGGCCATAACGG |
| *PCCB* | 7 (+ strand) | CAATGAGGATGTTACCCAGG |
| *ECHS1* | 2 (+ strand) | CCTGGTTGAGCTCGTCAATC |
| *ECHS1* | 3 (+ strand) | CCATTGACAGCAGCGATGACTGG |
| *ACADSB* | 6 (+ strand) | TTAGTAGATCGTGATACTCC |
| *ACADSB* | 6 (- strand) | ATCACGATCTACTAAGAAGG |
| *ACAD8* | 6 (+ strand) | GTGGTCATGTGCCGAACAGG |
| *ACADM* | 7 (+ strand) | AAGATGTGGATAACCAACGG |
| *ACADM* | 5 (+ strand) | GGGGTTCAGACTGCTATTGA |

**Table S2. Primers for PCR amplification and Sanger sequencing.**

| **Target gene** | **Exon (fragment size in bp)** | **Primer sequence** |
|---|---|---|
| *PCCB* | 5 (377bp) | Fw: 5’- TCGTGATGGGGAGTGAAAC -3’<br>Rev: 5’- GAACAGACACAATGCGGCAG -3’ |
| *PCCB* | 7 (306bp) | Fw: 5’- CCTCATTCAGCCACGTCACTA -3’<br>Rev: 5’- TAAAAACTGCCACCAGGCTCT -3’ |
| *ECHS1* | 2 (311bp) | Fw: 5’- CCCTTTTTGCCTTCTTGCTCC -3’<br>Rev: 5’- AGAGGCGCAAATCTGTCTGG -3’ |
| *ECHS1* | 3 (274bp) | Fw: 5’- CTGGTGTGCTCTGAAGTTCC -3’<br>Rev: 5’- CACGGTTCCTTCAACCACGA -3’ |
| *ACADSB* | 6 (378bp) | Fw: 5’- ATGTTGCAATTCTGAGTCCATTAG -3’<br>Rev: 5’- AATACTCTCCTGCGTGTGCTT -3’ |
| *ACAD8* | 6 (392bp) | Fw: 5’- TCCCAGACGCTGTGAGAACT -3’<br>Rev: 5’- TCCCCAGAGGCCTAGTGTTTA -3’ |
| *ACADM* | 7 (398bp) | Fw: 5’- ATTGTGTAACAGAACCTGGAGCA -3’<br>Rev: 5’- ACAGAGTGCTAGCCTTCCTA -3’ |
| *ACADM* | 5 (466bp) | Fw: 5’- TTCTTATTGTGCCAGCCAGAAC -3’<br>Rev: 5’- GGCCAGAGTTATCAAGCCTCG -3’ |

#### Figure S1

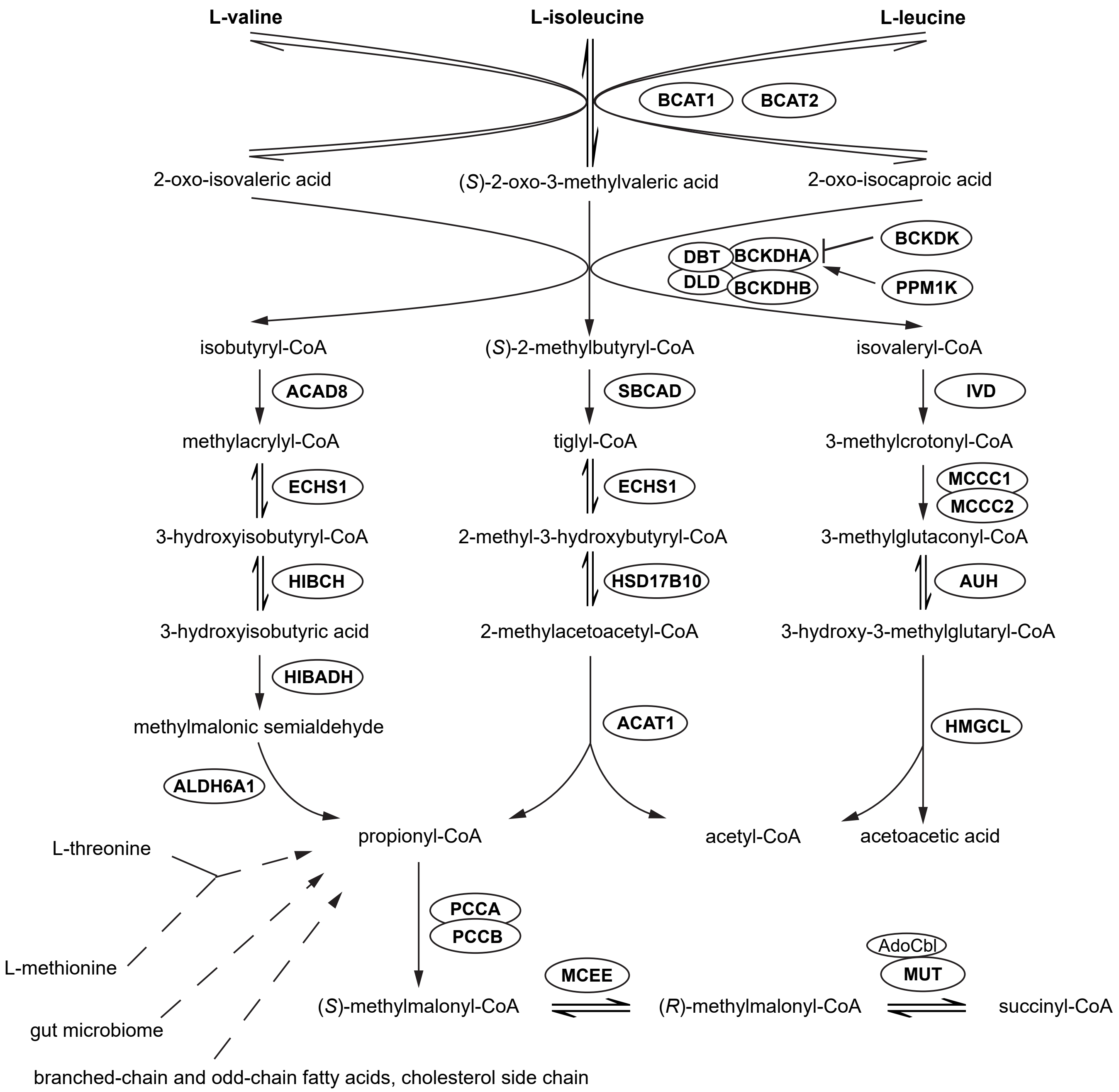

L-valine
L-isoleucine
L-leucine
BCAT1
BCAT2
2-oxo-isovaleric acid
(S)-2-oxo-3-methylvaleric acid
2-oxo-isocaproic acid
DBT
BCKDHA
DLD
BCKDHB
BCKDK
PPM1K
isobutyryl-CoA
(S)-2-methylbutyryl-CoA
isovaleryl-CoA
ACAD8
SBCAD
IVD
methylacrylyl-CoA
tiglyl-CoA
3-methylcrotonyl-CoA
ECHS1
ECHS1
MCCC1
MCCC2
3-hydroxyisobutyryl-CoA
2-methyl-3-hydroxybutyryl-CoA
3-methylglutaconyl-CoA
HIBCH
HSD17B10
AUH
3-hydroxyisobutyric acid
2-methylacetoacetyl-CoA
3-hydroxy-3-methylglutaryl-CoA
HIBADH
methylmalonic semialdehyde
ACAT1
HMGCL
ALDH6A1
L-threonine
propionyl-CoA
acetyl-CoA
acetoacetic acid
PCCA
PCCB
L-methionine
AdoCbl
MCEE
MUT
(S)-methylmalonyl-CoA
(R)-methylmalonyl-CoA
succinyl-CoA
gut microbiome
branched-chain and odd-chain fatty acids, cholesterol side chain

#### Figure S2

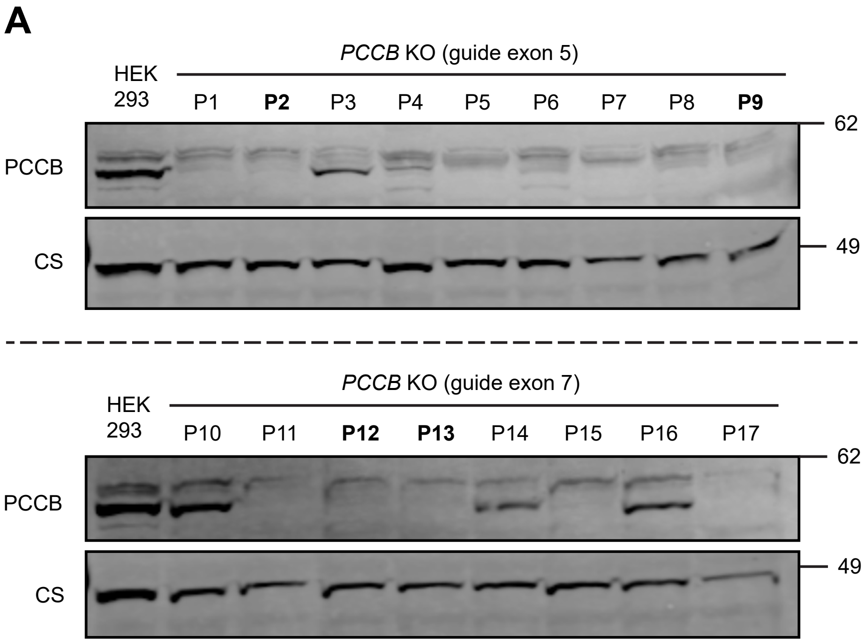
A
PCCB KO (guide exon 5)
HEK 293
P1
P2
P3
P4
P5
P6
P7
P8
P9
PCCB
CS
62
49
PCCB KO (guide exon 7)
HEK 293
P10
P11
P12
P13
P14
P15
P16
P17
PCCB
CS
62
49

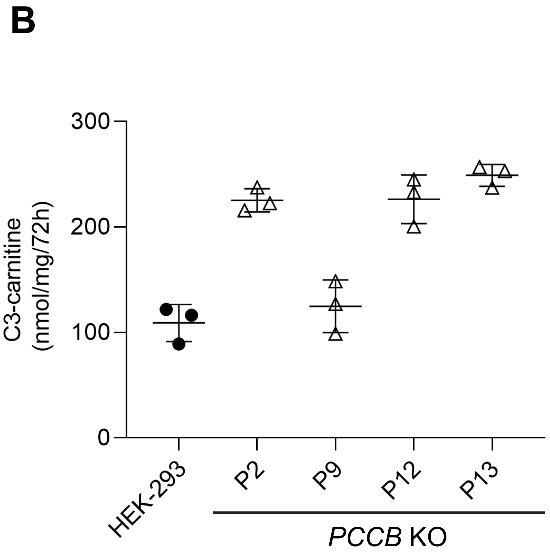
B
C3-carnitine (nmol/mg/72h)
300
200
100
0
HEK-293
P2
P9
P12
P13
PCCB KO

#### Figure S3

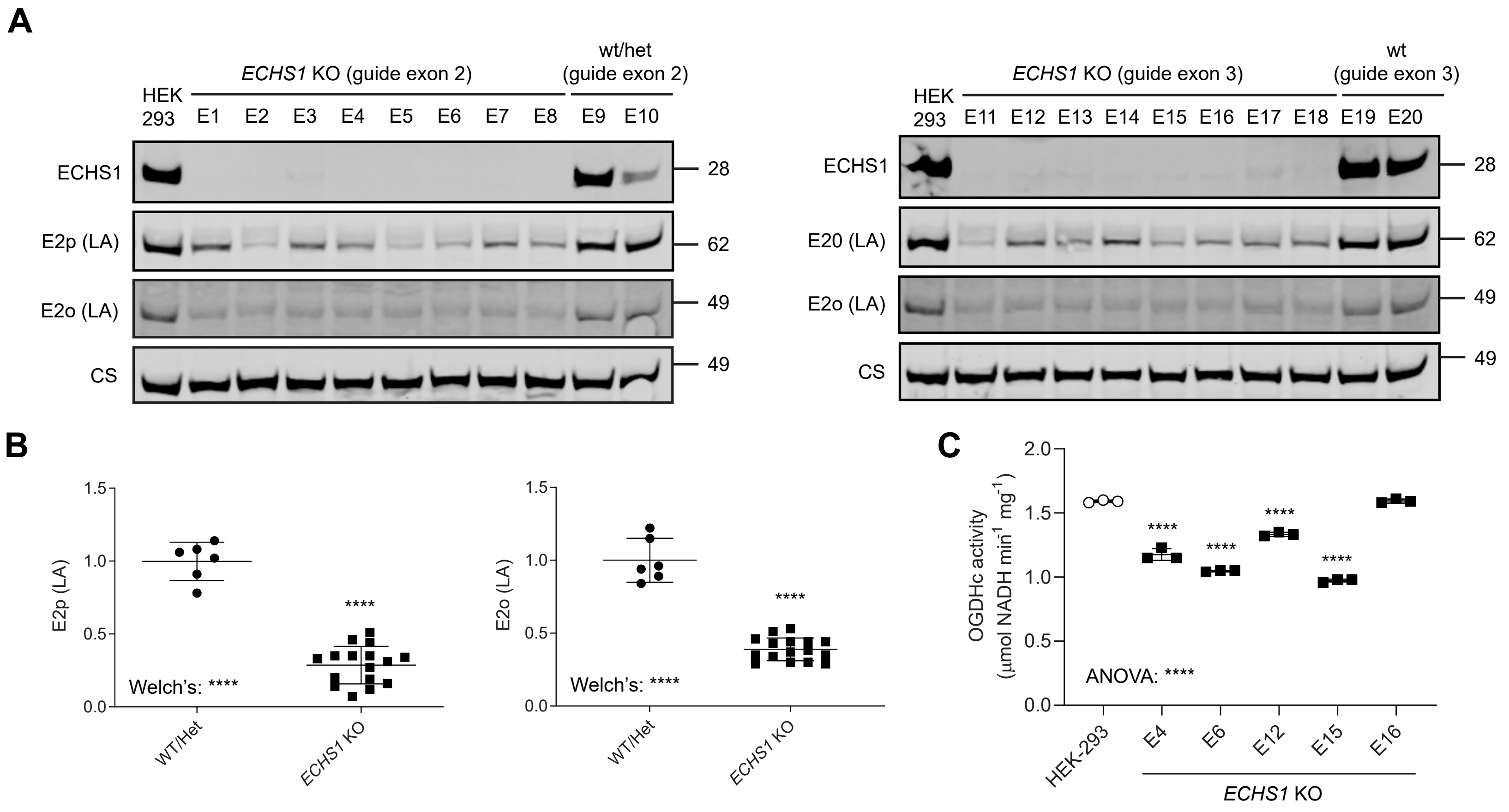

A
ECHS1 KO (guide exon 2)
wt/het (guide exon 2)
HEK 293
E1 E2 E3 E4 E5 E6 E7 E8 E9 E10
ECHS1
E2p (LA)
E2o (LA)
CS
28
62
49
49
ECHS1 KO (guide exon 3)
wt (guide exon 3)
HEK 293
E11 E12 E13 E14 E15 E16 E17 E18 E19 E20
ECHS1
E20 (LA)
E2o (LA)
CS
28
62
49
49
B
E2p (LA)
1.5
1.0
0.5
0.0
****
Welch's: ****
WT/Het
ECHS1 KO
E2o (LA)
1.5
1.0
0.5
0.0
****
Welch's: ****
WT/Het
ECHS1 KO
C
OGDHc activity (µmol NADH min-1 mg-1)
2.0
1.5
1.0
0.5
0.0
****
****
****
****
ANOVA: ****
HEK-293
E4
E6
E12
E15
E16
ECHS1 KO

### Figure S4

**A**

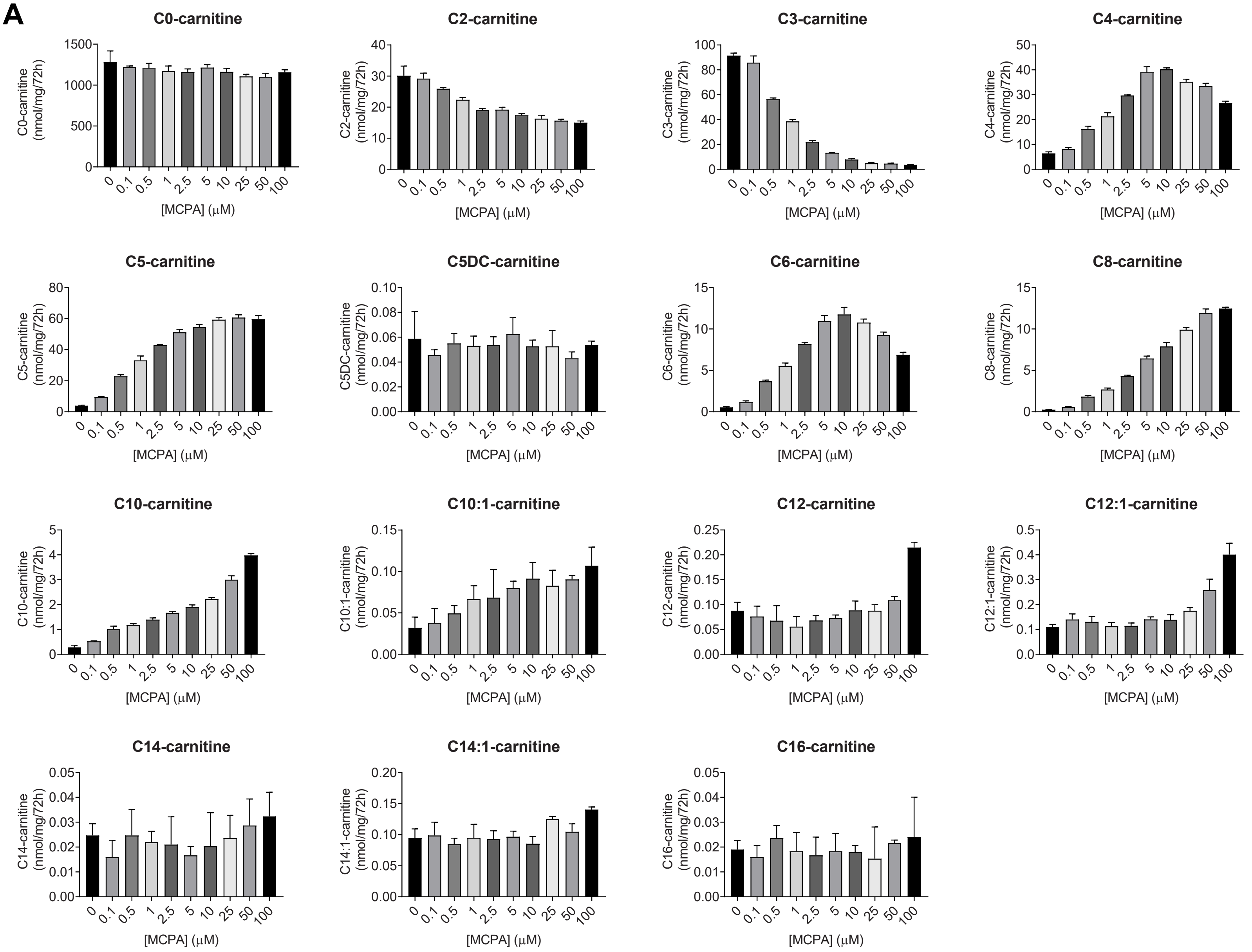


**B**

C4-carnitine isomers

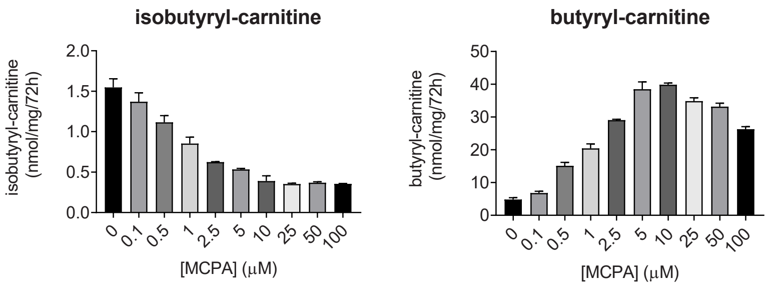


C5-carnitine isomers

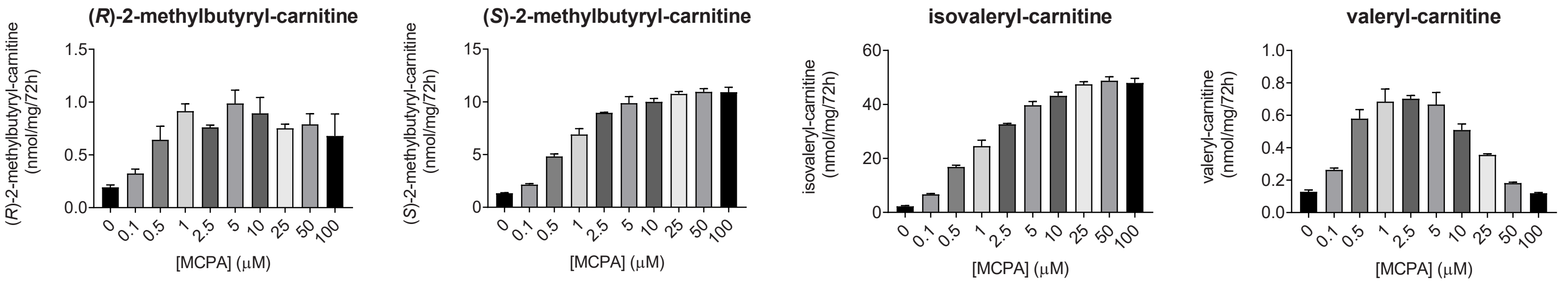

#### Figure S5

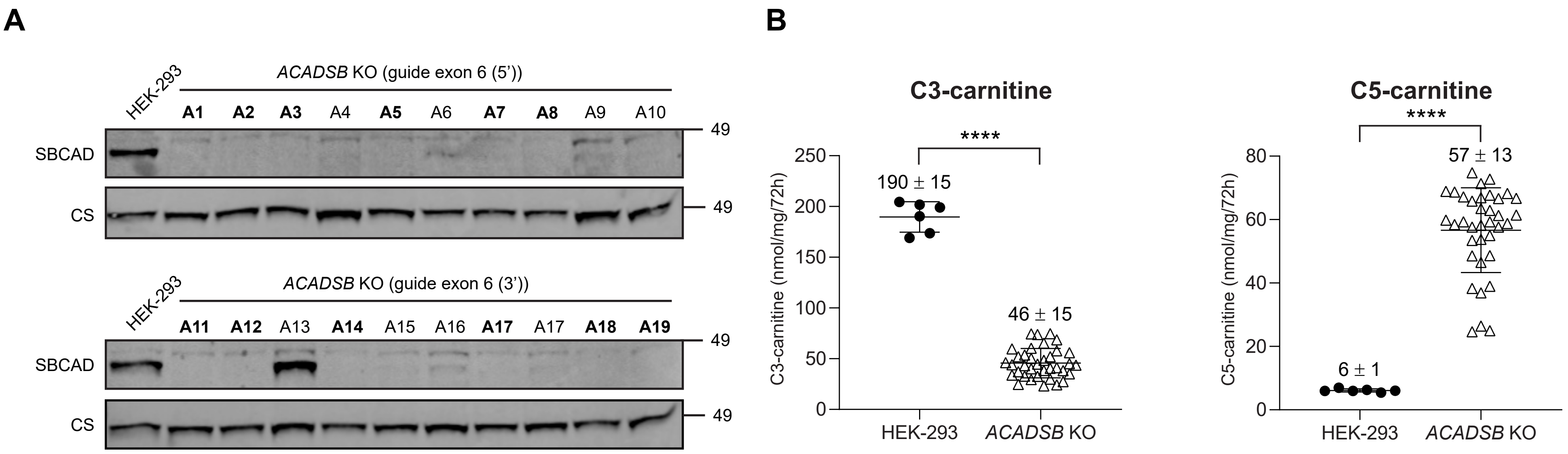
A
HEK-293
ACADSB KO (guide exon 6 (5'))
A1 A2 A3 A4 A5 A6 A7 A8 A9 A10
SBCAD
CS
49
HEK-293
ACADSB KO (guide exon 6 (3'))
A11 A12 A13 A14 A15 A16 A17 A17 A18 A19
SBCAD
CS
49
B
C3-carnitine
C3-carnitine (nmol/mg/72h)
****
190 ± 15
46 ± 15
0
50
100
150
200
250
HEK-293
ACADSB KO
C5-carnitine
C5-carnitine (nmol/mg/72h)
****
57 ± 13
6 ± 1
0
20
40
60
80
HEK-293
ACADSB KO

#### Figure S5

**C**

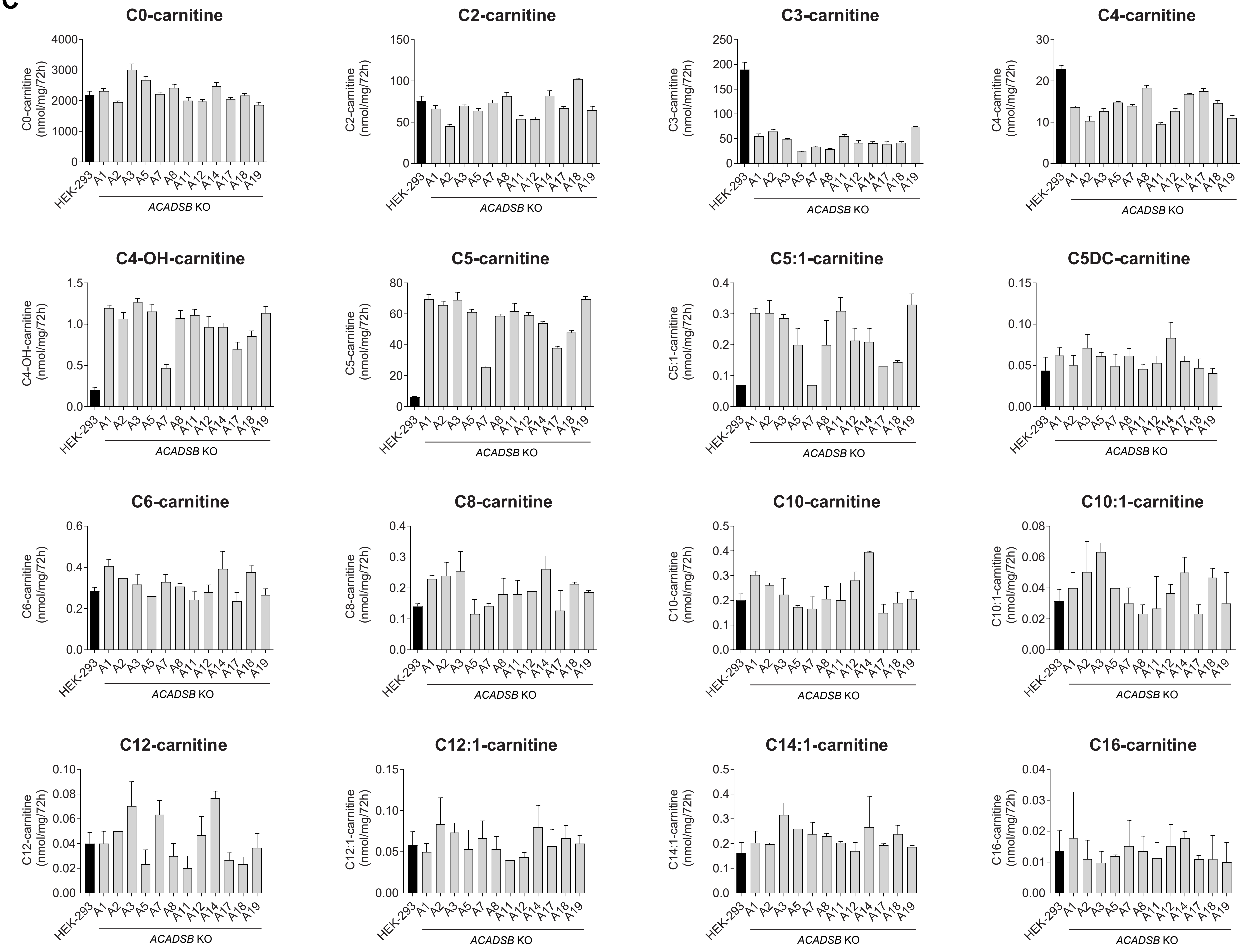
C0-carnitine
C0-carnitine (nmol/mg/72h)
C2-carnitine
C2-carnitine (nmol/mg/72h)
C3-carnitine
C3-carnitine (nmol/mg/72h)
C4-carnitine
C4-carnitine (nmol/mg/72h)
C4-OH-carnitine
C4-OH-carnitine (nmol/mg/72h)
C5-carnitine
C5-carnitine (nmol/mg/72h)
C5:1-carnitine
C5:1-carnitine (nmol/mg/72h)
C5DC-carnitine
C6-carnitine
C6-carnitine (nmol/mg/72h)
C8-carnitine
C8-carnitine (nmol/mg/72h)
C10-carnitine
C10-carnitine (nmol/mg/72h)
C10:1-carnitine
C10:1-carnitine (nmol/mg/72h)
C12-carnitine
C12-carnitine (nmol/mg/72h)
C12:1-carnitine
C12:1-carnitine (nmol/mg/72h)
C14:1-carnitine
C14:1-carnitine (nmol/mg/72h)
C16-carnitine
C16-carnitine (nmol/mg/72h)
HEK-293
A1
A2
A3
A5
A7
A8
A11
A12
A14
A17
A18
A19
ACADSB KO

#### Figure S6

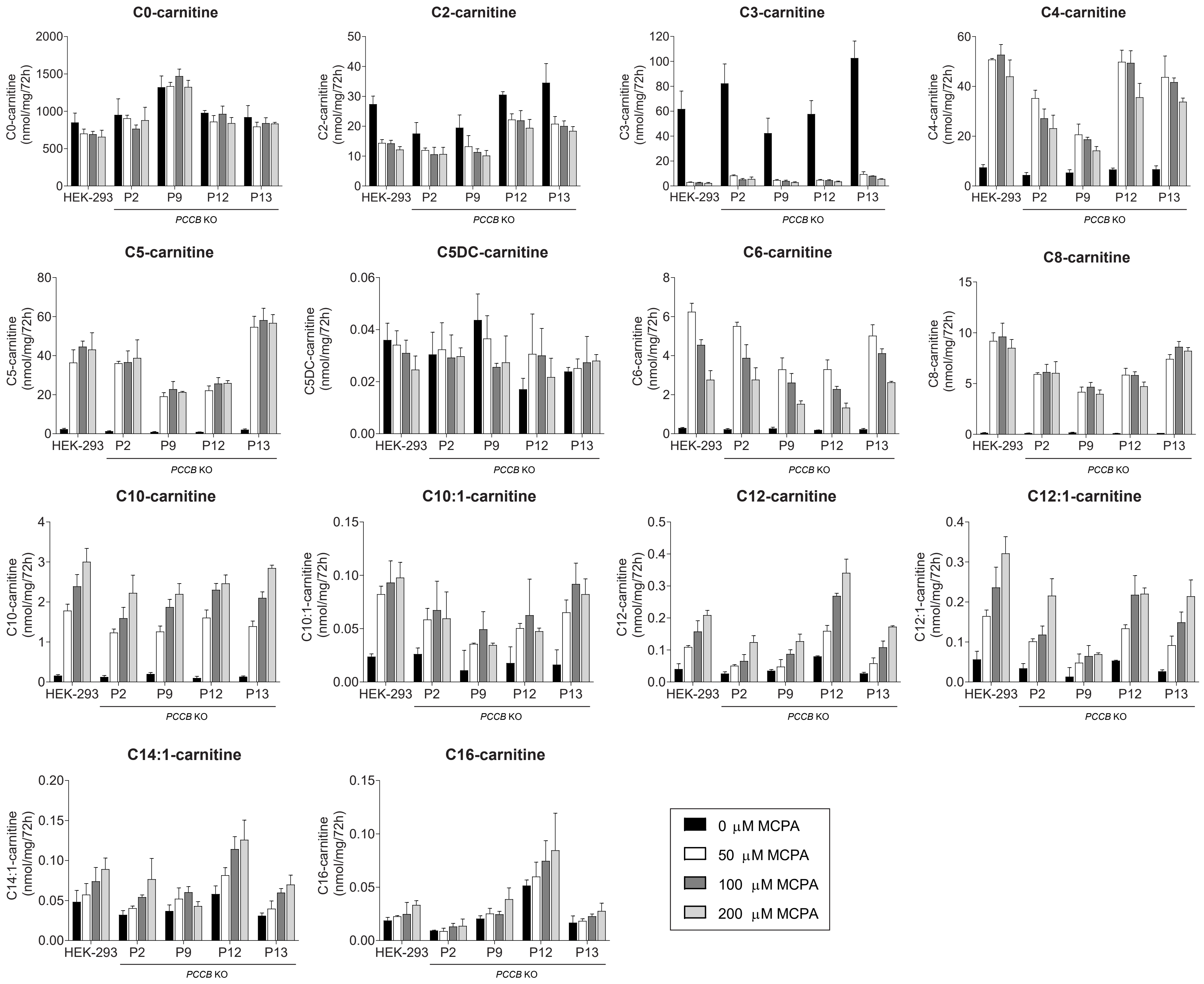
C0-carnitine
C2-carnitine
C3-carnitine
C4-carnitine
C5-carnitine
C5DC-carnitine
C6-carnitine
C8-carnitine
C10-carnitine
C10:1-carnitine
C12-carnitine
C12:1-carnitine
C14:1-carnitine
C16-carnitine
(nmol/mg/72h)
HEK-293
P2
P9
P12
P13
PCCB KO
0 μM MCPA
50 μM MCPA
100 μM MCPA
200 μM MCPA

### Figure S7

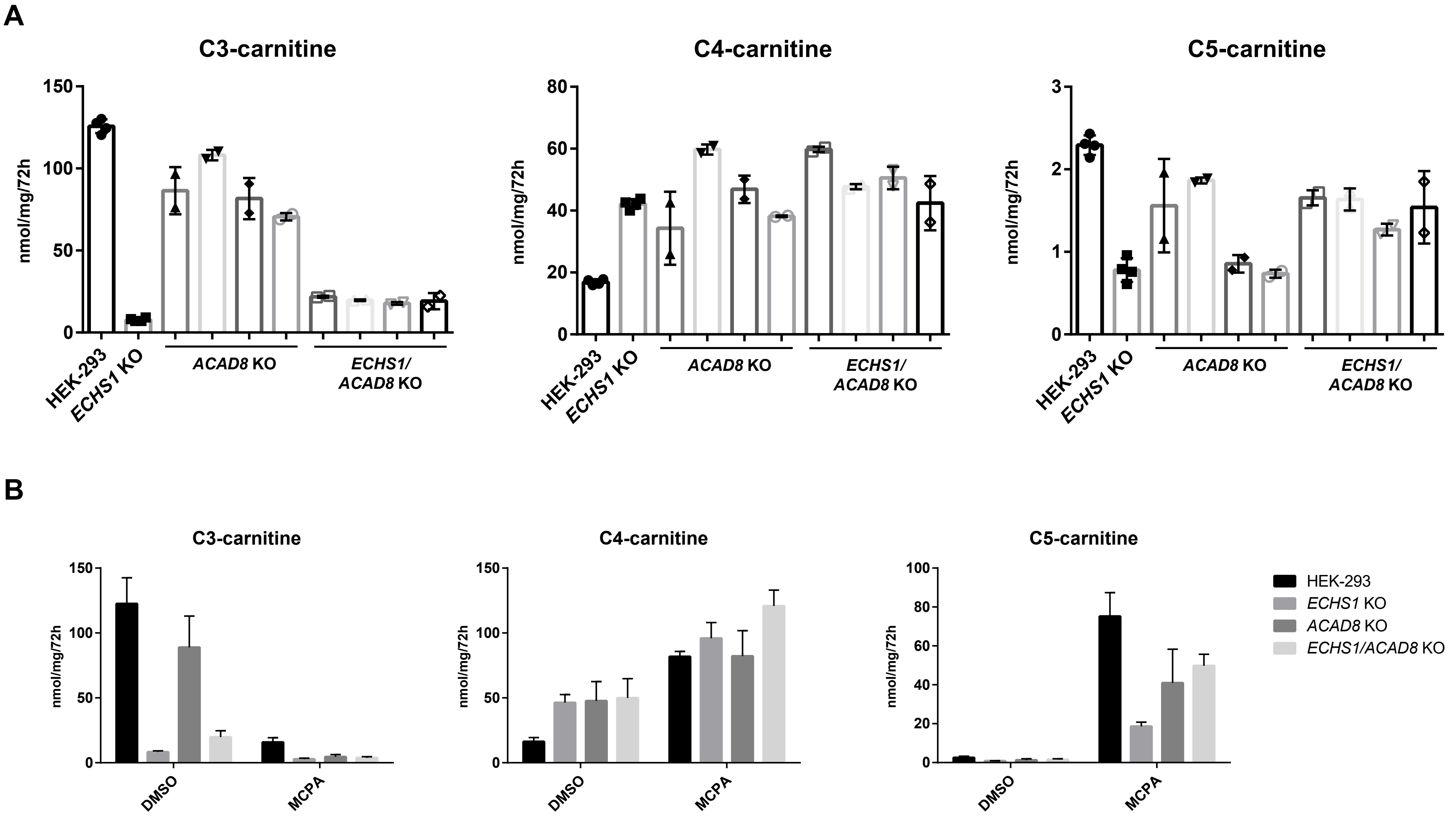
A
C3-carnitine
C4-carnitine
C5-carnitine
nmol/mg/72h
HEK-293
ECHS1 KO
ACAD8 KO
ECHS1/ ACAD8 KO
B
C3-carnitine
C4-carnitine
C5-carnitine
nmol/mg/72h
DMSO
MCPA
HEK-293
ECHS1 KO
ACAD8 KO
ECHS1/ACAD8 KO

#### Figure S8

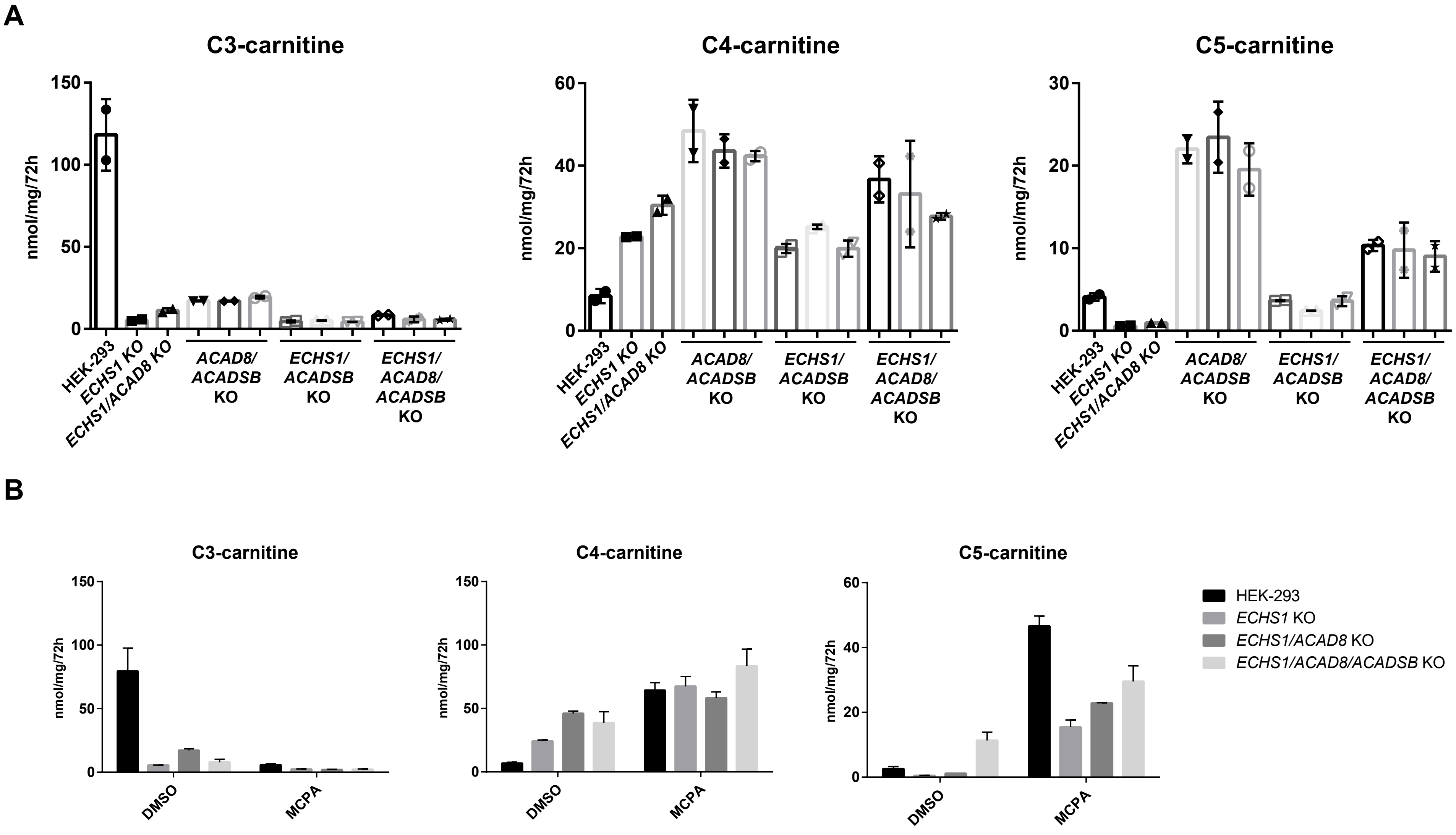

A
C3-carnitine
C4-carnitine
C5-carnitine
nmol/mg/72h
150
100
50
0
60
40
20
30
10
HEK-293
ECHS1 KO
ECHS1/ACAD8 KO
ACAD8/ ACADSB KO
ECHS1/ ACADSB KO
ECHS1/ ACAD8/ ACADSB KO
B
C3-carnitine
C4-carnitine
C5-carnitine
nmol/mg/72h
DMSO
MCPA
HEK-293
ECHS1 KO
ECHS1/ACAD8 KO
ECHS1/ACAD8/ACADSB KO

#### Figure S9

**A**

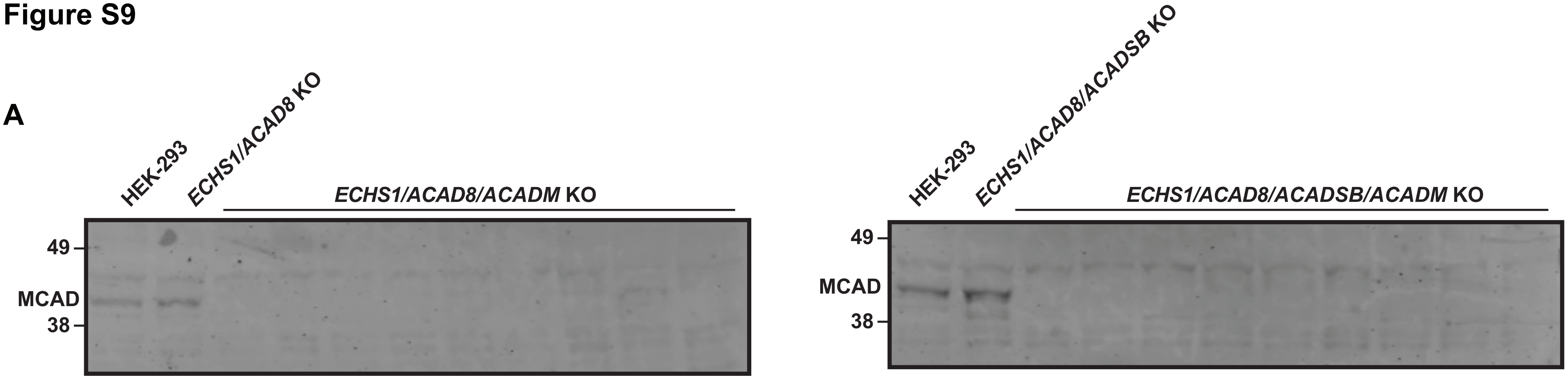
HEK-293
ECHS1/ACAD8 KO
ECHS1/ACAD8/ACADM KO
49
MCAD
38
HEK-293
ECHS1/ACAD8/ACADSB KO
ECHS1/ACAD8/ACADSB/ACADM KO
49
MCAD
38

**B**

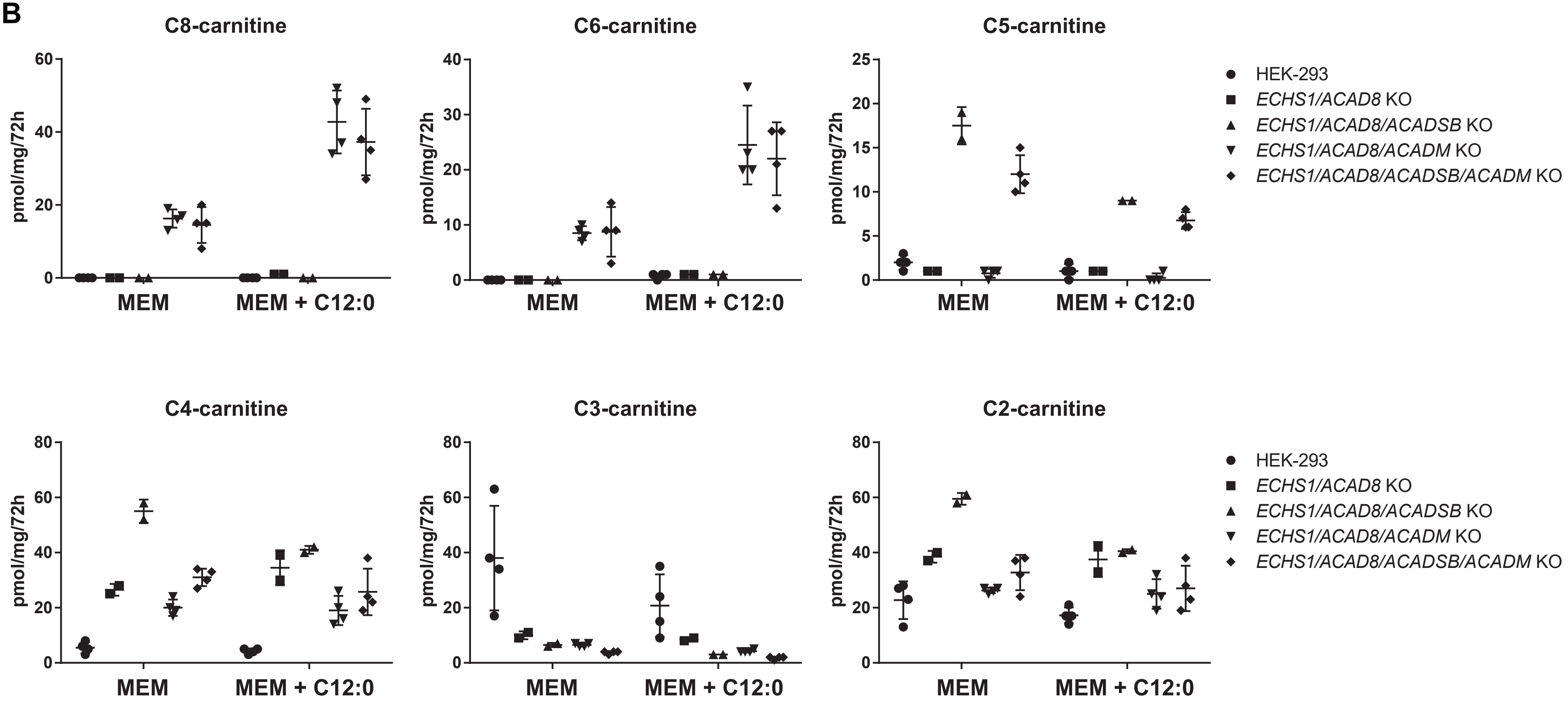
C8-carnitine
C6-carnitine
C5-carnitine
C4-carnitine
C3-carnitine
C2-carnitine
pmol/mg/72h
MEM
MEM + C12:0
HEK-293
ECHS1/ACAD8 KO
ECHS1/ACAD8/ACADSB KO
ECHS1/ACAD8/ACADM KO
ECHS1/ACAD8/ACADSB/ACADM KO
